# TGF-β family ligands exhibit distinct signaling dynamics that are driven by receptor localization

**DOI:** 10.1101/565416

**Authors:** Daniel S. J. Miller, Bernhard Schmierer, Caroline S. Hill

**Author notes:** Present address (Daniel S. J. Miller). Institute of Cancer Research, 15 Cotswold Road, Sutton, SM2 5NG, United Kingdom.

## Abstract

Growth factor-induced signal transduction pathways are tightly regulated at multiple points intracellularly, but how cells monitor levels of extracellular ligand and translate this information into appropriate downstream responses remains unclear. Understanding signaling dynamics is thus a key challenge in determining how cells respond to external cues. Here, we demonstrate that different TGF-β family ligands, namely Activin A and BMP4, signal with distinct dynamics, which differ profoundly from those of TGF-β itself. The distinct signaling dynamics are driven by differences in the localization and internalization of receptors for each ligand, which in turn determine the capability of cells to monitor levels of extracellular ligand. Using mathematical modeling, we demonstrate that the distinct receptor behaviors and signaling dynamics observed may be primarily driven by differences in ligand-receptor affinity. Furthermore, our results provide a clear rationale for the different mechanisms of pathway regulation found *in vivo* for each of these growth factors.

## Introduction

The transforming growth factor β (TGF-β) family of ligands plays diverse roles in embryonic development and adult tissue homeostasis, and moreover, their signaling is deregulated in a range of human diseases, including cancer (Massague, 2008, Pickup et al., 2017). The mammalian family consists of 33 members, which signal via the same conserved mechanism (Moses et al., 2016). Two classes of cell surface serine/threonine kinase receptors, termed type I and type II, recognize TGF-β family ligands. Ligand binding brings the receptors together, allowing the constitutively active kinase of the type II receptor to phosphorylate the type I receptor. This both activates the type I receptor, and provides a binding site for the intracellular effectors of the pathways, the SMADs (Heldin and Moustakas, 2016). The receptor-regulated SMADs (R-SMADs) become phosphorylated at their extreme C-termini by the type I receptor, which drives the formation of complexes with the common mediator SMAD, SMAD4. These complexes accumulate in the nucleus where they regulate the transcription of a battery of target genes in conjunction with specific co-factors. The TGF-β family has traditionally been split into two pathways, with the TGF-βs, NODAL and Activin leading to the phosphorylation of SMAD2/3, whereas the BMPs and some of the GDFs induce phosphorylation of SMAD1/5/9 (Schmierer and Hill, 2007). This, however, is a simplification, as some ligands, in particular TGF-β and Activin, can activate both signaling arms (Daly et al., 2008, Hatsell et al., 2015, Ramachandran et al., 2018).

TGF-β receptors are known to internalize in the absence and presence of ligand, and once activated, to signal from early endosomes (Di Guglielmo et al., 2003, He et al., 2015, Miller et al., 2018, Mitchell et al., 2004). A proportion of internalized receptors have been shown to recycle constitutively back to the cell surface, while the remainder are targeted for degradation (Le Roy and Wrana, 2005, Yakymovych et al., 2018). Although the mechanisms underlying the immediate cellular response to TGF-β family ligands is relatively well understood, the response to longer durations of ligand exposure, and the resulting dynamics of signaling, have been much less studied. All the mammalian TGF-β family ligands signal through just seven type I and five type II receptors, so the wide range of cell behaviors seen in response to different ligands are likely to involve additional levels of complexity, some of which will be at the level of signaling dynamics. Because cells are exposed to the continuous presence of TGF-β family ligands during embryonic development and in disease states (Hill, 2017, Miller and Hill, 2016, Schier and Talbot, 2005), as well as in the context of regenerative medicine (Pagliuca et al., 2014), it is crucial to understand how long-term exposure to ligands is regulated. This will be essential for identifying potential novel points of intervention in each pathway, both experimentally and for the development of therapeutic strategies. Moreover, as all TGF-β family ligands result in the phosphorylation of just two classes of R-SMAD, understanding whether particular ligands lead to different dynamic patterns of SMAD phosphorylation, and how these are regulated, is critical for our understanding of how these pathways evolved and diverged.

We have previously shown that in response to the continuous presence of TGF-β, cells enter a refractory state where they no longer respond to acute TGF-β stimulation. This is due to the rapid depletion of receptors from the cell surface in response to ligand (Vizan et al., 2013). This means that intracellular signaling downstream of TGF-β (as read out, for example, by levels of phosphorylated R-SMADs) is not proportional to the duration of signaling, neither is it sensitive to the presence of ligand antagonists in the extracellular milieu. This type of behavior would clearly be incompatible with the ability of ligands like BMPs, NODAL and Activin to act as morphogens that signal over many cell diameters in the context of embryonic development and tissue homeostasis (Langdon and Mullins, 2011, Hedger and de Kretser, 2013). We thus postulated that these other TGF-β family ligands might respond to prolonged ligand exposure in a different manner to TGF-β.

We set out to directly test this hypothesis by fully characterizing the response of cells to prolonged Activin and BMP4 stimulation. Our results show that in contrast to TGF-β, cells integrate their response to BMP4 and Activin over time, and do not enter a refractory state when stimulated with these ligands. Moreover, we observe an oscillatory SMAD1/5 phosphorylation in response to BMP4 stimulation, which we show is driven by the transient expression of the I-SMADs, SMAD6 and SMAD7, which leads to a temporary depletion of receptors from the cell surface. By combining our experimental insights with mathematical modeling we can explain these distinct behaviors of Activin, BMP4 and TGF-β by differences in trafficking of their cognate receptors, and differential affinities of ligands for their receptors. This in turn may explain the distinct functional roles these ligands play *in vivo*.

## Results

### BMP4 and Activin exhibit distinct patterns of signaling dynamics

We have previously shown that when cells are stimulated with TGF-β, SMAD2 phosphorylation peaks after 1 hr, before attenuating to lower levels. After an initial acute response, cells are refractory to further acute stimulation due to an almost complete depletion of receptors from the cell surface (Vizan et al., 2013). To understand whether this was a common feature of all TGF-β family ligands, we characterized the response of cells to other members of the TGF-β family, namely Activin A and BMP4, and compared and contrasted them with each other and with TGF-β. For the Activin responses we have predominantly used the P19 mouse teratoma cell line, as SMAD2 is robustly phosphorylated in response to Activin in this cell line (Coda et al., 2017). Activin signaling in these cells is mediated by ACVR1B as the type I receptor, and either ACVR2A or ACVR2B as the type II receptors, as demonstrated by the abrogation of signaling when these receptors are knocked down by siRNA (Figure 1 – figure supplement 1). These cells also produce and secrete the TGF-β family ligands NODAL and GDF3, resulting in a relatively high level of basal level of SMAD2 phosphorylation (Coda et al., 2017). To characterize the BMP4 responses we have predominantly used the human breast cancer cell line, MDA-MB-231 and the mouse fibroblast cell line NIH-3T3, both of which induce robust SMAD1/5 phosphorylation in response to BMP4. In addition, we have used HaCaTs, the cell line we previously used to characterize TGF-β signaling dynamics (Vizan et al., 2013).

In response to continuous stimulation with BMP4, SMAD1/5 phosphorylation in MDA-MB-231 cells peaks after 1 hr, then drops down to a lower level after 4 hr, before increasing back up to its maximal level after 8 hr of stimulation (Figure 1A). This is strikingly different to the dynamics of signaling seen in response to TGF-β, where chronic exposure of cells to ligand leads to signal attenuation resulting in a low level of SMAD2 phosphorylation (Vizan et al., 2013). A similar single oscillation is evident when NIH-3T3 cells (Figure 1 – figure supplement 2A) or human keratinocyte HaCaT cells (Figure 1 – figure supplement 2B) are stimulated with BMP4, although NIH-3T3s reach their low point of signaling after 2 hr of stimulation, rather than 4 hr, and in neither of these cell types does the signal return to the maximal level, as in does in the MDA-MB-231 line. The long term response to Activin is different. P19 cells stimulated with Activin exhibit maximal levels of PSMAD2 after 1 hr, which modestly attenuates down to the basal level over the next 24 hr (Figure 1B). Basal PSMAD2 is completely abolished by overnight incubation with the type I receptor inhibitor SB-431542 (Inman et al., 2002a) (Figure 1B). P19s can also be induced by Activin from the SB-431542-inhibited baseline, and in this case, show a very sustained response, due to the autocrine production of NODAL and GDF3 (Coda et al., 2017). In HaCaTs, in contrast, the baseline of PSMAD2 is low and the Activin response is more transient, likely because HaCaTs do not exhibit autocrine signaling (Figure 1C)

**Figure 1.**
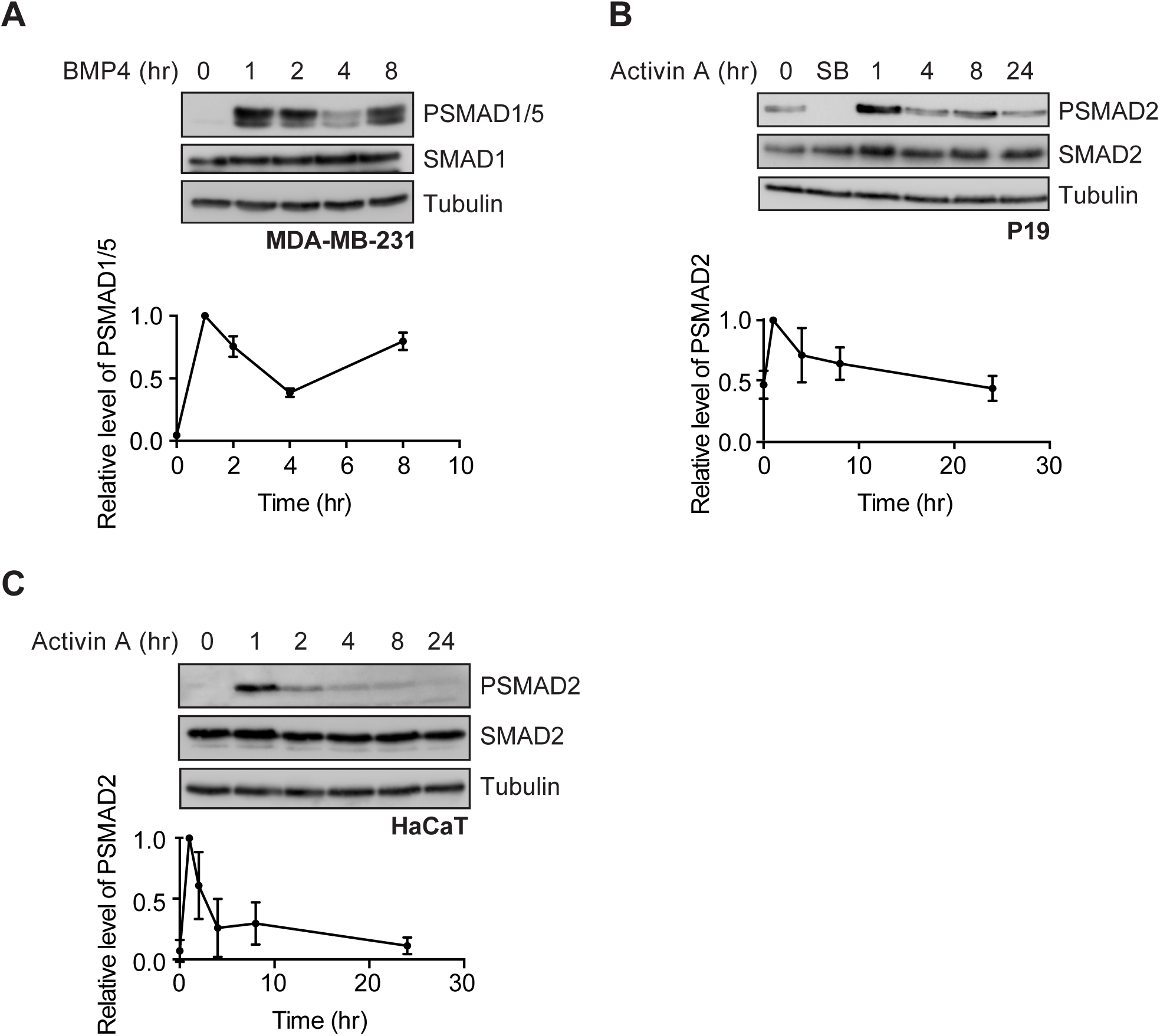
BMP4 and Activin signal with distinct dynamics. **(A)** MDA-MB-231 cells were treated with BMP4 for the times indicated. **(B** and **C)** P19 cells (**B**) or HaCaT cells (**C**) were treated with Activin A for the times indicated or SB-431542 (SB) overnight. Western blotting for PSMAD1/5, SMAD1, PSMAD2, SMAD2/3 and Tubulin as a loading control was performed. Quantifications are the normalized means and standard deviations (SDs) of densitometry measurements from three independent experiments.

### Activin and BMP4 signaling is integrated over time

We next sought to determine whether signaling by Activin and BMP4 is integrated over time after stimulation, and compared the behaviors with that TGF-β. Cells were therefore stimulated for increasing periods of time with Activin, BMP4, or TGF-β, and then chased for the remainder of the 1 hr with saturating doses of the natural ligand antagonists, Follistatin (Nakamura et al., 1990) or Noggin (Zimmerman et al., 1996) for Activin and BMP4 respectively, or, in the case of TGF-β, the neutralizing antibody 1D11 (Nam et al., 2008) (Figure 2A). All cells were harvested together at the 1 hr time point. TGF-β induced a maximal PSMAD2 response after just 5 mins of exposure to ligand (Figure 2B), which we have previously demonstrated is due to the rapid depletion from the cell surface of the type II TGF-β receptor TGFBR2 within this time frame, so that little to no new signaling is induced over the remainder of the first hour of signaling (Vizan et al., 2013). In contrast, the cellular response to Activin is integrated over the first hour of signaling, with a greater induction of PSMAD2 resulting from longer exposure to ligand (Figure 2C and D). A similar pattern was observed with SMAD1/5 phosphorylation resulting from BMP4 stimulation in MDA-MB-231 cells (Figure 2E) and HaCaT cells (Figure 2F). We conclude that cells continuously monitor the presence of BMP4 and Activin in their extracellular environment, such that the R-SMAD phosphorylation observed after 1 hr in response to BMP and Activin is an integration of all of the signaling that has occurred in the first hour. This behavior is distinct from that of TGF-β, where the SMAD phosphorylation seen after 1 hr of stimulation is the result of the first 5 minutes of ligand exposure.

**Figure 2.**
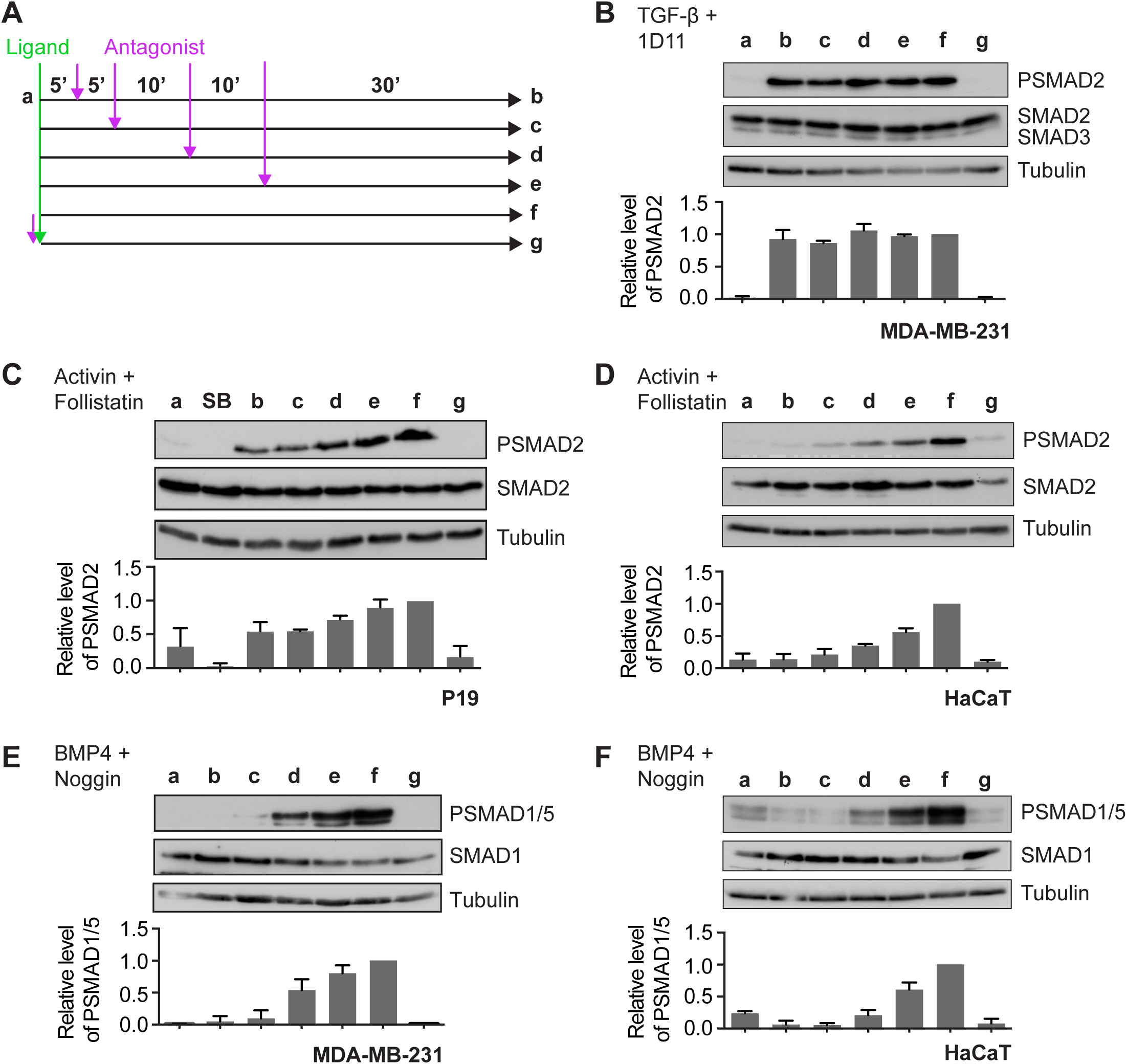
Activin and BMP4 signals are integrated over time, whilst TGF-β signals are not. **(A)** Experimental scheme. Cells were untreated (a), or treated with ligand for 5 (b), 10 (c), 20 (d), 30 (e), or 60 (f) minutes, followed by the cognate ligand antagonist for the remainder of 60 min. To ensure that inhibitors were working as expected, cells were pre-treated with inhibitor for 5 mins, followed by ligand for 60 mins (g). **(B)** MDA-MB-231 cells were treated as in (A) with TGF-β and the blocking antibody, 1D11. **(C)** P19 cells were treated as in (A) with Activin and Follistatin, and additionally overnight with SB-431542 (SB). **(D)** HaCaT cells were treated as in (A) with Activin and Follistatin. **(E)** MDA-MB-231 cells were treated as in (A) with BMP4 and Noggin. **(F)** HaCaT cells were treated as in (A) with BMP4 and Noggin. Western blotting for PSMAD1/5, SMAD1, PSMAD2, SMAD2/3 and Tubulin as a loading control was performed. Quantifications are the normalized means and SDs of densitometry measurements from three independent experiments.

### Stimulation with Activin and BMP4 does not induce refractory behavior

We have previously shown that cells enter a refractory state in response to TGF-β treatment, where they are unable to respond to acute stimulation with the same ligand. To determine whether the same state is induced in response to Activin and BMP4, cells were stimulated with these ligands for 1 hr, followed by ligand antagonists for 2 hr to reduce R-SMAD phosphorylation levels down to basal. The ligand antagonists were then washed out and cells re-stimulated with ligand for 1 hr. The efficacy of the ligand antagonists and their wash-out was confirmed (Figure 3A and B). For both BMP4 (Figure 3A) and Activin (Figure 3B), re-stimulation to maximal PSMAD levels was observed after just 2 hr treatment with ligand antagonists, indicating that cells do not enter a refractory state in response to these ligands. This contrasts with the behavior of cells stimulated with TGF-β. In this case, where cells take 12–24 hr after the removal of external ligand to recover the ability to fully respond again to ligand (Vizan et al., 2013).

**Figure 3.**
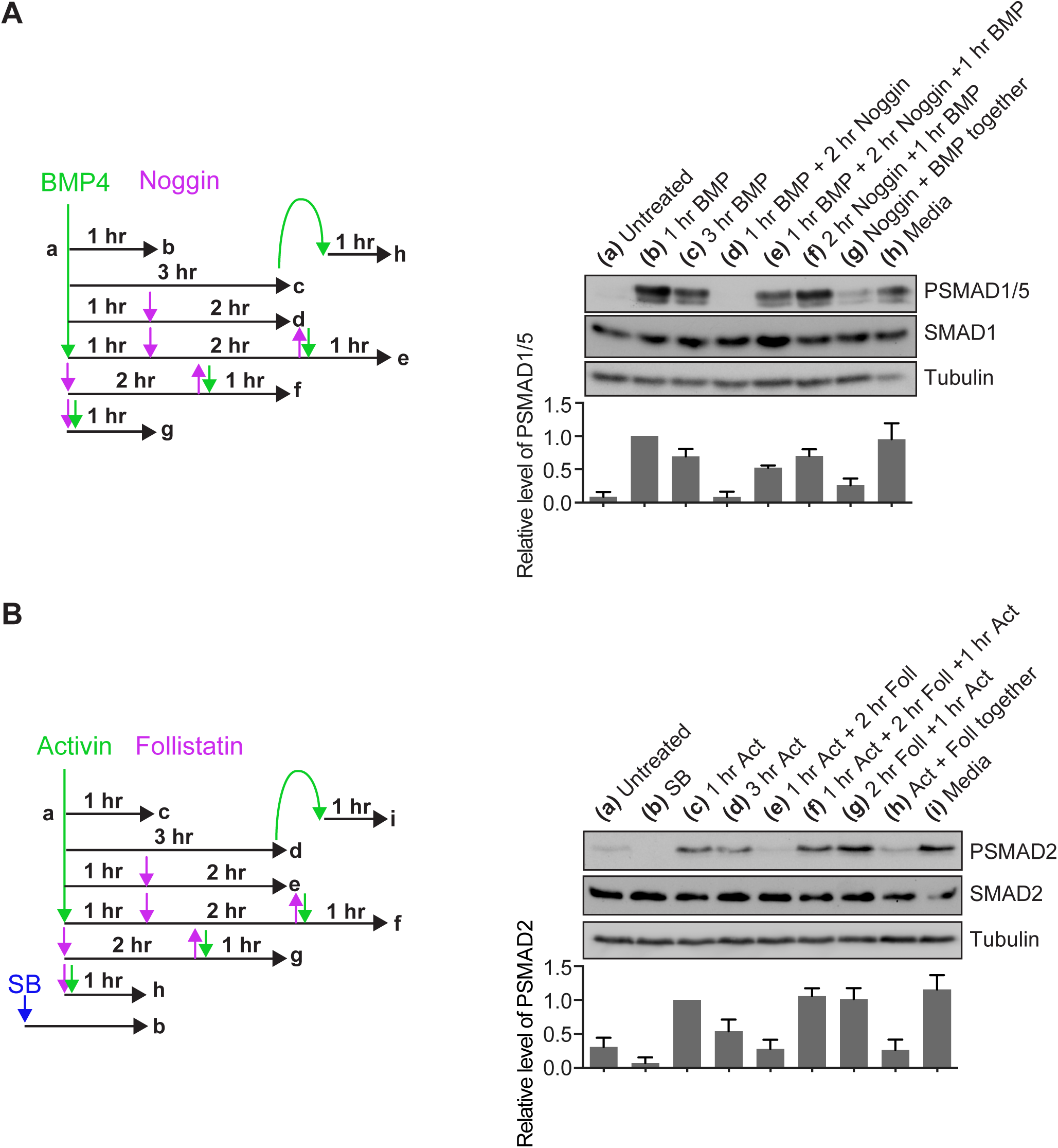
BMP4 and Activin do not induce refractory behavior. **(A)** Left, a schematic of experimental set-up. NIH-3T3 cells were untreated (a) or treated with BMP4 for 1 hr (b) or 3 hr (c). After 1 hr of BMP4 stimulation, signal was brought down to baseline with Noggin for 2 hr (d), which was then washed out and cells re-stimulated with BMP4 for 1 hr (e). The efficacy of Noggin washout was confirmed (f), as was its inhibitory ability by adding the ligand and antagonist simultaneously (g). To confirm that BMP4 was not depleted from the media in the time period of these experiments, cells were stimulated with BMP4 for 3 hr, then the media transferred to naïve cells for 1 hr (h). Western blotting for PSMAD1/5, SMAD1 and Tubulin as a loading control was performed. Quantifications are the normalized means and SDs of densitometry measurements from three independent experiments. **(B)** Left, a schematic of experimental set-up. P19 cells were untreated (a), treated overnight with SB-431542 (b) or treated with Activin for 1 hr (c) or 3 hr (d). After 1 hr of Activin stimulation, signal was brought down to baseline with Follistatin for 2 hr (e), which was then washed out and cells re-stimulated with Activin for 1 hr (f). The efficacy of Follistatin washout was confirmed (g), as was its inhibitory ability (g). To confirm that Activin was not depleted from the media in the time period of these experiments, cells were stimulated with Activin for 3 hr, then the media transferred to naïve cells for 1 hr (i). Western blotting for PSMAD2, SMAD2 and Tubulin as a loading control was performed. Quantifications are the normalized means and SDs of densitometry measurements from three independent experiments.

### The distinct signaling dynamics of TGF-β, Activin and BMP4 are not explained by the intracellular lifetimes of their receptors

TGF-β family receptors can signal from internal cellular compartments (Itoh et al., 2002), and we have shown that the lifetime of receptors in these compartments is likely to be an important factor for regulating the dynamics of signaling (Vizan et al., 2013). We therefore determined whether the distinct signaling dynamics observed in response to each ligand could be driven by the duration for which activated receptors signal from internal compartments.

To address this, cells were stimulated for 1 hr with TGF-β, BMP4 or Activin, then chased over a time course of 8 hr with the cognate ligand antagonists 1D11, Noggin and Follistatin respectively, and the levels of R-SMAD phosphorylation were assayed (Figure 4). Because there is no new signaling induced by the activation of receptors with external ligands once antagonists are added, any on-going PSMAD signal must arise from the combined activities of the receptors signaling from internalized compartments, and cellular R-SMAD phosphatases. To control for the latter, the decay in R-SMAD phosphorylation due to the action of R-SMAD phosphatases was assayed directly by chasing stimulated cells with receptor kinase inhibitors SB-431542 (for TGF-β and Activin) or LDN-193189 (for BMP4; Cuny et al., 2008) over the same time course. By comparing the decay in signal seen with ligand antagonists versus receptor kinase inhibitors, the duration of signaling from internal compartments can be determined. In the presence of the kinase inhibitors, maximal R-SMAD dephosphorylation occurred within around 30 mins in all cases (Figure 4B–D), with a half-life of approximately 15 mins, measured by fitting an exponential decay curve to the data. In contrast, in the presence of ligand antagonists, the signal in response to TGF-β decayed with a half-life of approximately 52 mins, the signal from BMP4 in approximately 44 mins and that from Activin in approximately 42 mins (Figure 4B–D). Thus, signaling persists for around 2 hr in all cases, suggesting that receptors signal from endosomes for approximately 90 mins, with no obvious differences seen between the different ligands.

**Figure 4.**
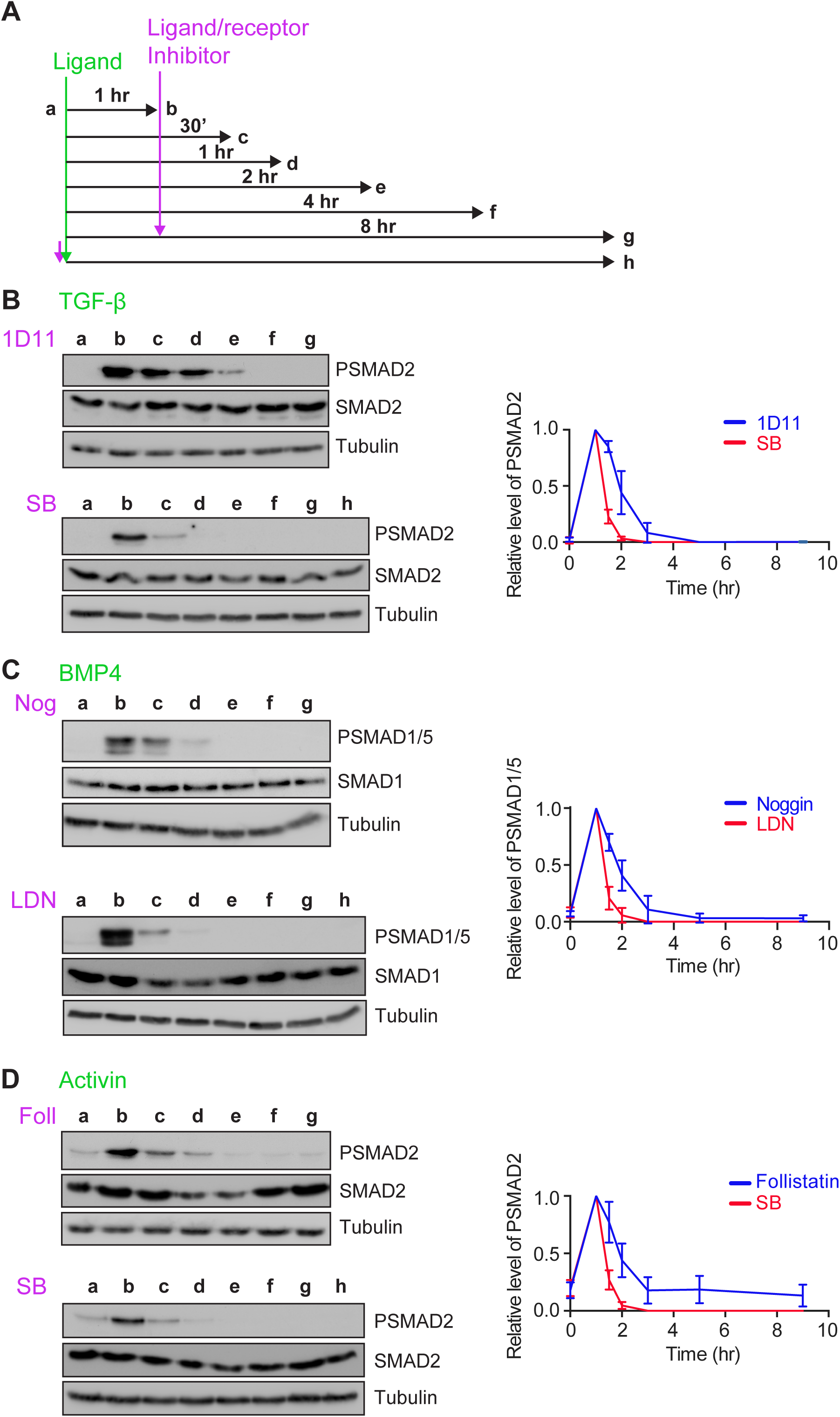
The distinct TGF-β, Activin and BMP4 signaling dynamics are not explained by the intracellular lifetimes of their receptors. **(A)** Experimental scheme. Cells were untreated (a), or treated for 1 hr with ligand (b), then with ligand antagonist or receptor kinase inhibitors for 30 mins (c), 1 hr (d), 2 hr (e), 4 hr (f) or 8 hr (g), or with ligand and receptor kinase inhibitor together for 8 hr (h). **(B)** MDA-MB-231 cells were treated as in (A) with TGF-β, 1D11 or SB-431542 (SB). **(C)** MDA-MB-231 cells were treated as in (A) with BMP4, Noggin (Nog) or LDN-193189 (LDN). **(D)** P19 cells were treated as in (A) with Activin A, Follistatin (Foll) or SB-431542 (SB). Western blotting for PSMAD1/5, SMAD1, PSMAD2, SMAD2/3 and Tubulin as a loading control was performed. Quantifications are the normalized means and SDs of densitometry measurements from three independent experiments.

### Receptor trafficking behaviors drive distinct signaling dynamics for the different ligands

We reasoned that differences in signaling dynamics could be driven by differences in the behavior of the receptors for each ligand. Antibodies for Western blot were validated against one of the BMP4 type II receptors, BMPR2 (Daly et al., 2008), the Activin type I receptor ACVR1B (formerly known as ALK4) and ACVR2B (Tsuchida et al., 2009). In all cases, PNGase treatment, which removes N-linked sugars, resulted in an increased mobility of the receptors, and siRNA knockdown was used to confirm the specificity of the antibodies and the identity of the correct band (Figure 5 – figure supplement 1A–C).

We next determined the half-life of each receptor species to which we had antibodies. Cycloheximide chase time courses were performed, which showed that BMPR2 has a half-life of approximately 4 hr (Figure 5 – figure supplement 1D), and ACVR1B approximately 1 hr (Figure 5 – figure supplement 1E). ACVR2B was not noticeably degraded at all over the time course in either P19s (Figure 5 – figure supplement 1F) or HaCaTs (Figure 5 – figure supplement 1G), indicating that it has a much longer half-life than the other receptors tested. The half-lives of BMPR2 and ACVR1B are of the same order as those previously calculated for the TGF-β receptors (~ 2 hr for TGFBR2 and ~4 hr for TGFBR1; Vizan et al., 2013).

We previously showed that TGFBR1 and TGFBR2 become rapidly depleted from the surface of cells in response to TGF-β stimulation (Vizan et al., 2013). We therefore wanted to know whether BMP and Activin stimulation similarly drives receptor depletion, and used surface biotinylation assays on cells treated with BMP4 or Activin to test this. In MDA-MB-231 cells, BMPR2 was depleted from the cell surface after 2 hr of BMP4 treatment, before re-accumulating at later time points (Figure 5A). Although receptors re-accumulated, they did not appear to fully reach their level in unstimulated cells. Receptor depletion and re-accumulation occurs with similar dynamics to the oscillation in PSMAD1/5 levels seen in response to signal. Despite the transient depletion of BMPR2, in response to long-term stimulation, it remains present at the cell surface. This explains why cells do not become refractory to further acute stimulation after treatment with BMP4.

**Figure 5.**
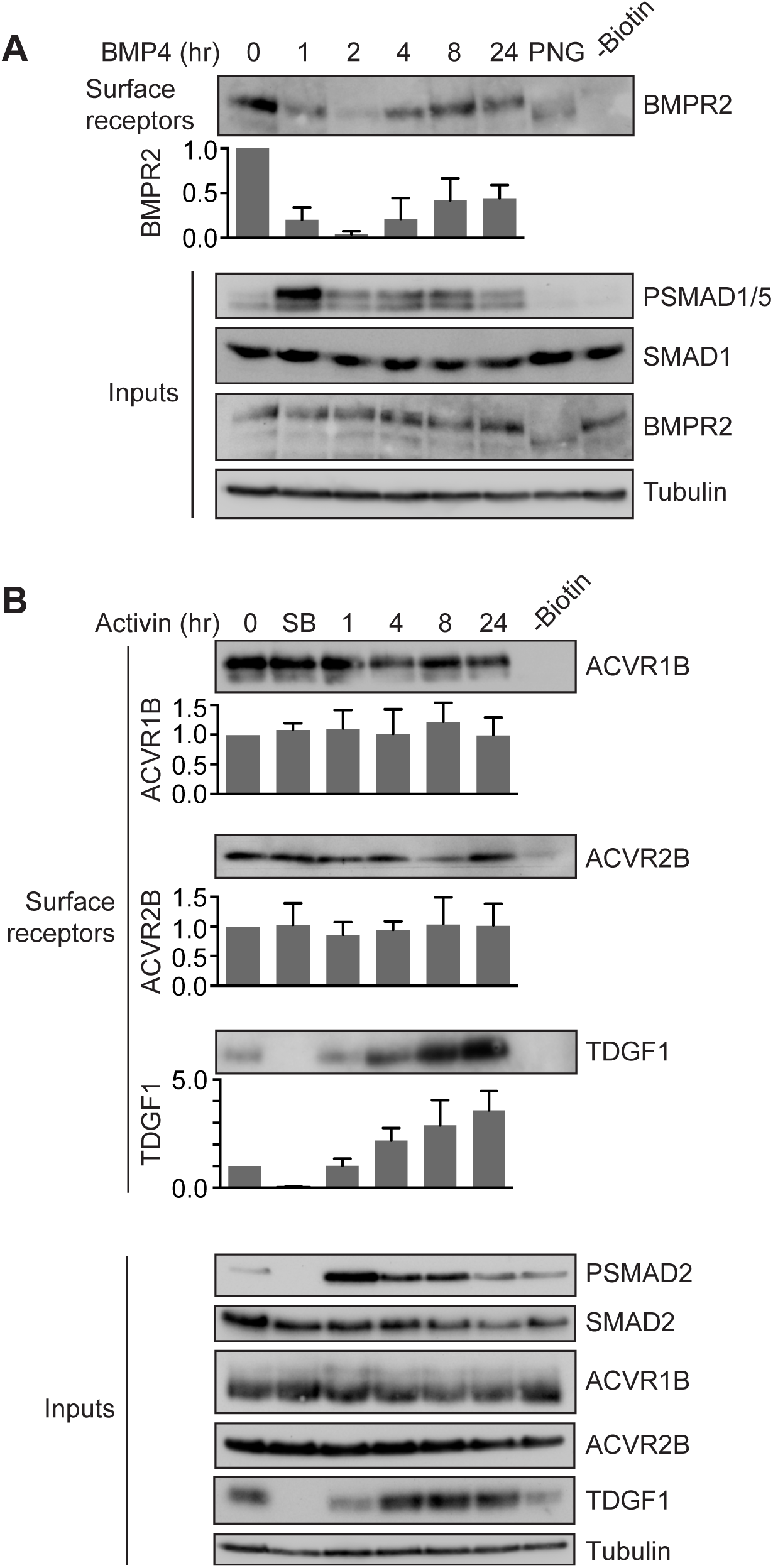
BMP4 and Activin drive distinct receptor trafficking behaviors. **(A)** MDA-MB-231 cells were treated with BMP4 for the times indicated. **(B)** P19 cells were treated with Activin for the times indicated or SB-431542 overnight (SB). Whole cell extracts were Western blotted for BMPR2, PSMAD1/5, SMAD1, ACVR1B, ACVR2B, TDGF1, PSMAD2, SMAD2/3 with Tubulin as a loading control (Inputs). Surface biotinylation assays were performed to isolate surface receptor populations, which were Western blotted for BMPR2, ACVR1B, ACVR2B and TDGF1. For the lanes marked -Biotin unstimulated cell extracts were treated identically to the other samples, but without the addition of Biotin. In A, the lane marked PNG corresponds to a 0 time point where the sample was treated with PNGase to remove N-linked sugars from the receptors prior to gel electrophoresis. Quantifications are the normalized means and SDs of densitometry measurements from three independent experiments, relative to the levels in untreated cells.

By contrast, using P19 cells, we could show that neither ACVR1B nor ACVR2B deplete from the cell surface in response to Activin or in the presence of the receptor inhibitor, SB-431542 (Figure 5B). As a control for visualization of a cell surface protein, whose levels change in response to signal, we assessed the cell surface levels of the NODAL/GDF co-receptor TDGF1, whose expression is up-regulated in response to Activin signaling. TDGF1 robustly accumulated in response to Activin both at the cell surface and in whole cell lysates (Figure 5B). Again, the constant presence of Activin receptors at the cell surface during ligand stimulation explains why cells do not enter a refractory state after an acute Activin induction. Cells thus remain competent to respond to acute doses of ligand in their extracellular environment, even after an initial stimulation with Activin.

### The oscillatory response to BMP4 depends on the continuous presence of BMP4 in the extracellular milieu and requires new protein synthesis

Stimulation with BMP4 leads to an oscillatory PSMAD1/5 response driven by receptor depletion and re-accumulation. This oscillatory behavior is visible in multiple cell lines from different species, including NIH-3T3s, MDA-MB-231s and HaCaTs, although they show slightly different time points at which PSMAD1/5 reaches its nadir, and recover to different extents (Figure 1, Figure 1 – figure supplement 2). Because NIH-3T3 cells exhibited the most robust oscillation, they were used for subsequent experiments. To determine if the second wave of signaling after the dip in PSMAD1/5 is a result of new receptor activation at the cell surface or a second wave of signaling from internalized receptors, cells were stimulated for 1 hr with BMP4, which was subsequently washed out (Figure 6 – figure supplement 1A) or neutralized with Noggin (Figure 6 – figure supplement 1B). In both cases, no second wave of signaling was seen, indicating that the continuous presence of BMP4 in the media is necessary for the second increase in PSMAD1/5 observed after an initial decrease.

One possible explanation for these oscillatory dynamics is that another TGF-β family ligand, such as TGF-β itself, could be playing a role, possibly as a feedback target of the pathway that could be negatively regulating SMAD1/5 phosphorylation (Gronroos et al., 2012). To exclude this possibility, at least for a large subset of ligands that signal through SMAD2/3, BMP4 time courses were performed in the presence and absence of the TGF-β/Activin/Nodal receptor inhibitor, SB-431542 (Figure 6 – figure supplement 1C). However, no differences in PSMAD1/5 dynamics in response to BMP4 were seen in the presence or absence of SB-431542, ruling out such a feedback mechanism.

We also investigated whether protein synthesis was required for the oscillatory behavior. Time courses of BMP4 treatment were performed in the presence or absence of the translation inhibitor, cycloheximide. In the presence of cycloheximide, no oscillation in PSMAD1/5 levels was observed and levels remained high throughout the time course (Figure 6 – figure supplement 2A). This indicates that a negative regulator of the pathway must be expressed in response to signaling, and that this factor is responsible for oscillatory PSMAD1/5 dynamics. To confirm this, time courses of BMP4 treatment were performed in the presence of the transcriptional inhibitor Actinomycin D (Figure 6 – figure supplement 2B). Again, the dip in PSMAD1/5 levels seen in control cells is abrogated in the absence of new transcription.

### The oscillatory response to BMP4 requires the inhibitory SMADs, SMAD6 and SMAD7

Two of the most likely candidates to be feedback inhibitors of BMP signaling are the inhibitory SMADs (I-SMADs), SMAD6 and SMAD7. Both I-SMADs have long been known to be targets of BMP signaling (Takase et al., 1998) and are negative regulators of the pathway. Several mechanisms for their inhibitory activity have been proposed, including interfering with SMAD complex formation (Hata et al., 1998), inhibiting R-SMAD phosphorylation (Hayashi et al., 1997, Nakao et al., 1997, Imamura et al., 1997), targeting receptors for degradation (Ebisawa et al., 2001, Kavsak et al., 2000) or blocking the DNA binding and transcriptional activity of the SMADs (Lin et al., 2003). In NIH-3T3s, qPCR revealed that in response to BMP4 stimulation, *Smad6* and *Smad7* mRNAs are both induced in a transient manner that is the exact inverse of the PSMAD1/5 signal for *Smad6*, and in phase with PSMAD1/5 signal for *Smad7* (Figure 6A). siRNA-mediated knockdown of *Smad6* and *Smad7* together abrogated the oscillation in PSMAD1/5 levels seen with control, non-targeting (NT) siRNAs (Figure 6B). Individual siRNA pools against *Smad6* and *Smad7* both abolish oscillations in PSMAD1/5, although knockdown of *Smad7* leads to a weaker PSMAD1/5 response and a reduction in total SMAD1 levels (Figure 6 – figure supplement 2C).

**Figure 6.**
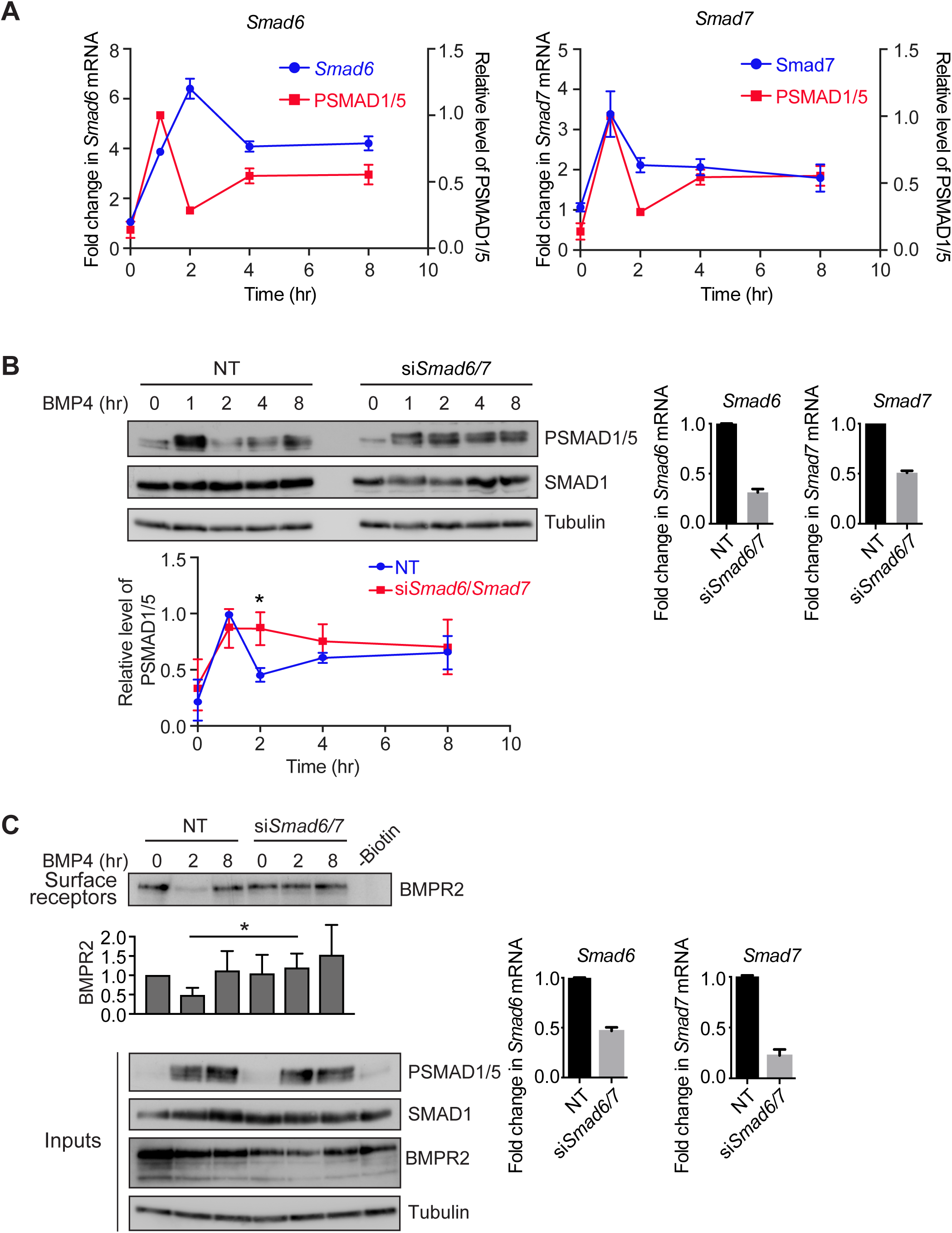
SMAD6 and SMAD7 are required for the oscillatory signaling response to BMP4. **(A)** NIH-3T3 cells were treated with BMP4 for the times indicated. Levels of *Smad6* and *Smad7* mRNA were assayed by qPCR. Shown are the normalized means and SDs from three independent experiments, expressed as fold change in mRNA level relative to untreated cells, overlaid with SMAD1/5 phosphorylation data from Figure 2 – figure supplement 2. **(B)** NIH-3T3 cells were transfected with non-targeting control siRNAs (NT) or siRNA SMARTpools targeting *Smad6* and *Smad7*, and were then treated with BMP4 for the times indicated. Western blotting for PSMAD1/5, SMAD1 and Tubulin was performed. Quantifications are the normalized means and SDs of densitometry measurements from three independent experiments. * indicates p<0.05. The extent of knockdown was determined by qPCR. Shown are the normalized means and SDs from three independent experiments, expressed as fold change in mRNA level relative to NT controls. (**C**) NIH-3T3 cells were transfected with non-targeting control siRNAs (NT) or siRNA SMARTpools targeting *Smad6* and *Smad7*, and were then treated with BMP4 for the times indicated. A biotinylation assay was performed to isolate surface receptor populations, which were Western blotted for BMPR2. Input cell lysates were also Western blotted for BMPR2, PSMAD1/5, SMAD1 and Tubulin as a loading control. For the lane marked -Biotin unstimulated cell extracts were treated identically to the other samples, but without the addition of Biotin. Quantifications are the normalized means and SDs of densitometry measurements from three independent experiments, relative to the levels in untreated cells. * indicates p<0.05. The extent of knockdown was determined by qPCR. Shown are the normalized means and SDs from three independent experiments, expressed as fold change in mRNA level relative to NT controls.

To confirm that these results apply across cell lines from different species, the dynamics of expression of *SMAD6* and *SMAD7* in response to BMP4 in MDA-MB-231 cells were also examined. *SMAD6* is induced after 2 hr of BMP4 stimulation and stays elevated over the duration of an 8-hr time course, while *SMAD7* shows a transient peak of expression after 2 hr, then declines down to a lower level (Figure 6 – figure supplement 3A). Knockdown of *SMAD6* and *SMAD7* together in MDA-MB-231 cells abrogates the PSMAD1/5 oscillation in a similar way to that observed in NIH-3T3 cells (Figure 6 – figure supplement 3B), indicating that this mechanism is conserved across species.

### SMAD6 and 7 are required for the transient depletion of BMPR2 from the cell surface

SMAD6 and SMAD7 have been described to target TGF-β superfamily receptors for degradation (Ebisawa et al., 2001, Goto et al., 2007, Kavsak et al., 2000). We therefore reasoned that the transient peak in their expression in response to BMP4 could be responsible for the transient depletion of BMP receptors from the cell surface, leading to the subsequent dip in SMAD1/5 phosphorylation. To test this, surface biotinylation assays were performed in NIH-3T3 cells transfected with either control NT siRNAs or siRNAs against *Smad6* and *Smad7*. The BMPR2 receptor also transiently depletes and re-accumulates in this cell line in response to BMP4 stimulation, indicating that this mechanism is conserved across species (Figure 6C). With knockdown of SMAD6 and SMAD7, BMPR2 was no longer transiently depleted from the cell surface in response to BMP4, but remained at high levels throughout the time course, indicating that a failure to deplete receptors from the cell surface in the absence of SMAD6 and SMAD7 underlies the lack of oscillation in SMAD1/5 phosphorylation in this condition (Figure 6C).

### Using mathematical modeling to find the key parameters that dictate specific signaling dynamics

Finally, we used mathematical modeling to obtain clues as to key parameters that might explain the distinct signaling dynamics of the different ligands. We previously built a mathematical model of the TGF-β pathway that simulated the refractory behavior of the TGF-β ligand (Vizan et al., 2013). Using this model as a starting point, we used our experimental findings, as well as the published literature, to determine whether, by changing some key parameters, we could simulate the signaling dynamics of Activin and BMP4 that we observe experimentally.

A striking difference between TGF-β itself and the other TGF-β family ligands is that TGF-β binds its receptors cooperatively, whilst there is no evidence for cooperativity in receptor binding for BMP4 and Activin (Hinck, 2012, Groppe et al., 2008). This likely explains the higher affinity measured for TGF-β1 and TGF-β3 for their receptors (*K*_d_ = 5–30 pM) (De Crescenzo et al., 2003, Massague, 1990), compared with the lower affinities measured for BMP4 and Activin with their cognate receptors (*K*_d_ = 110 pM for BMP4 and 100–380 pM for Activin) (Attisano et al., 1992, Luyten et al., 1994).

Starting first with the Activin pathway, we used our model to investigate whether lowering the affinity of Activin for its receptors would result in the distinct behaviors we have measured for Activin signaling versus TGF-β signaling. We found that implementing a *K*_d_ of 365 pM for Activin, and making minor adjustments to several other parameters (see Methods section) resulted in the model converting from simulating the characteristic behaviors of TGF-β signaling to those of Activin. The modified model faithfully captures the long-term Activin dynamics both in cells with no basal signaling, like HaCaTs, or with basal signaling, like P19s (Figure 7A). The simulations also reproduced the observed integration of signaling over time (Figure 7B), the behavior of the pathway when receptors are inhibited with a small molecule inhibitor, or when ligand is neutralised with Follistatin (Figure 7C), and also the ability of the pathway to be re-stimulated after ligand removal (Figure 7D).

**Figure 7.**
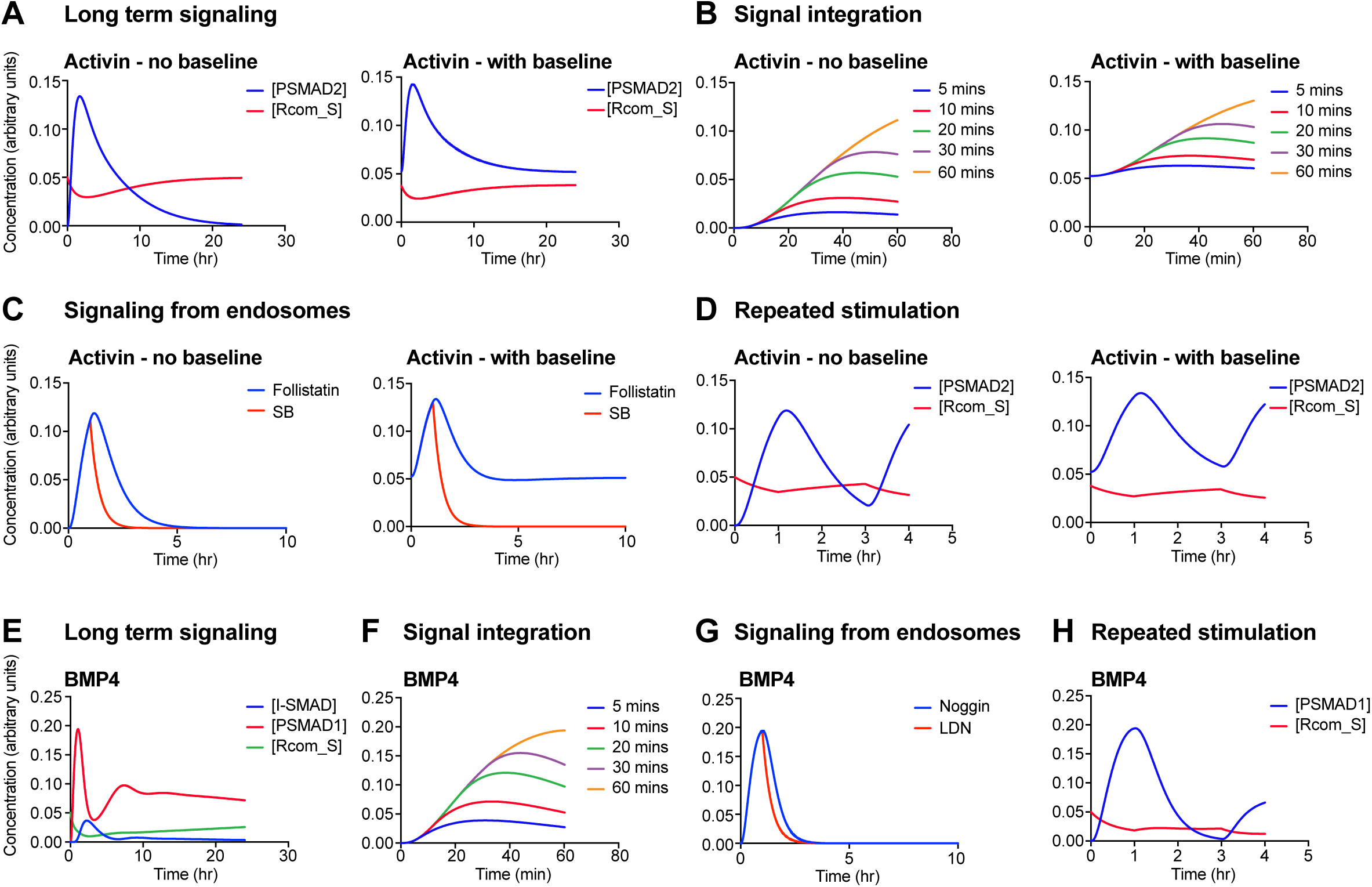
Mathematical models of the Activin and BMP pathways can simulate the experimentally-observed behaviors of these ligands. **(A–D)** The mathematical model was used to simulate the response of cells to Activin. In all cases, responses in cells with no baseline (e.g. HaCaTs) are shown on the left and responses in a cell line that has a basal level of PSMAD2 signaling (e.g. P19 cells) are shown on the right. **(A)** Simulation of a long-term Activin response; compare with experimental results in Figure 1C (HaCaTs) or Figure 1B (P19s). **(B)** Simulation of the signal integration experiments; compare with Figure 2D and Figure 2C respectively. **(C)** Simulation of the experiment shown in Figure 4D, which shows that signaling occurs from intracellular compartments, presumed to be endosomes. **(D)** Simulation of repeated Activin stimulation; compare Figure 3B. **(E–H)** Equivalent simulations were performed for the BMP4 responses. Compare **(E)** with Figure 1A; **(F)** with Figure 2E; **(G)** with Figure 4C and **(H)** with Figure 3A. In all cases concentrations of the indicated species are plotted in arbitrary units. In **(B)** and **(C)**, PSMAD2 concentration is plotted, and in **(F)** and **(G)**, PSMAD1 concentration is plotted.

BMP4 signaling dynamics are similar to Activin’s in the long term, but additionally show oscillatory behavior in the short term. We have shown that SMAD6 and SMAD7 are required for the oscillation, likely due to their role in inducing activated receptor degradation (Ebisawa et al., 2001, Kavsak et al., 2000). Their effect is transient, because expression of *Smad6* and *Smad7* in response to BMP4 is transient (Figure 6A). We implemented a *K*_d_ of 365 pM for BMP4 binding to its receptors, and additionally included the induction of SMAD6/7 by nuclear PSMAD1–SMAD4 complexes. This was implemented with an RNA intermediate and a non-linear dependency of *Smad6/7* expression on activatory PSMAD1–SMAD4 complexes. The SMAD6/7 is then assumed to act on the stability of activated receptors (see Methods section for the parameters and details of the modeling). This model captured all the main behaviors of BMP signaling that we observe experimentally, including the oscillation, signal integration over time, the behavior of the pathway when receptors are inhibited, or when ligand is neutralised with Noggin, and also the ability of the pathway to be re-stimulated after ligand removal (Figure 7E–H).

## Discussion

### Receptor trafficking and degradation dictates signaling dynamics for different TGF-β family ligands

In both physiological and pathological contexts *in vivo*, cells are frequently exposed to extracellular ligands for prolonged periods, yet little is currently understood about how cells respond to sustained ligand exposure, or about how signaling dynamics are modulated over time. In this study we have addressed these questions for members of the TGF-β family of ligands. We have shown that the signaling dynamics differ considerably between Activin, BMP4 and TGF-β and that they are dependent on the localization and behavior of cell surface receptors. In contrast to the behavior of cells treated with TGF-β, cells monitor the presence of Activin and BMP4 in the extracellular milieu during signaling, and as a result, signaling is integrated over time. Cells also do not enter a refractory state after an acute stimulation with Activin and BMP4, as they do in response to TGF-β. However, while continuous Activin stimulation leads to fairly stable SMAD2/3 phosphorylation in P19 cells, due to the continuous presence of receptors at the cell surface and autocrine signaling, BMP4 stimulation in a number of different cell lines leads to a transient depletion of the receptors from the cell surface due to the transient up-regulation of the I-SMADs, SMAD6 and SMAD7. This in turn results in an oscillatory signaling response to BMP4, where the response as read out by R-SMAD phosphorylation transiently dips and then recovers.

We therefore propose a model where the dynamics of signaling observed in response to different ligands of the TGF-β superfamily are determined by the localization and trafficking of cell surface receptors, specifically their rates of internalization from the cell surface and degradation, and their rates of renewal by recycling and/or new synthesis. At steady state prior to ligand induction, for all receptors, the rate of renewal matches the rate of depletion (Figure 8A). For TGF-β, ligand addition increases the rate of receptor internalization and degradation, so receptors become depleted from the cell surface and signaling attenuates (Figure 8B) (Vizan et al., 2013). For Activin, upon ligand addition, depletion is matched by renewal, such that receptors are not depleted from the cell surface. Moreover, the response to ligand is integrated until maximal R-SMAD phosphorylation is reached, and cells do not become refractory to acute stimulation (Figure 8C). For BMP4, receptor behavior over the first hour and in the longer term is similar to Activin, but a transient peak of SMAD6 and SMAD7 expression means that the rate of depletion and/or degradation is greater than the rate of renewal, leading to a transient dip in SMAD1 phosphorylation (Figure 8D).

**Figure 8.**
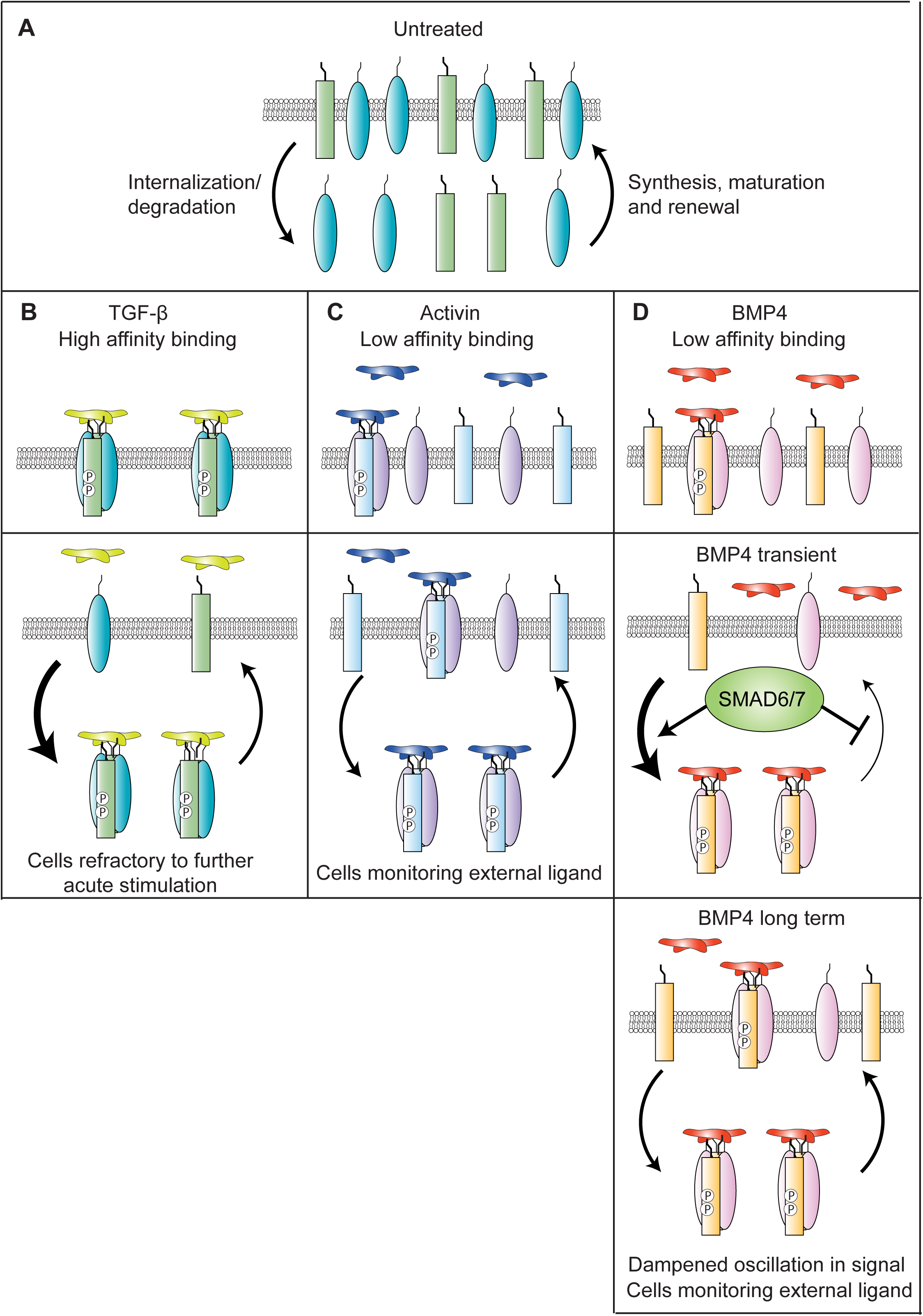
TGF-β family signaling dynamics are determined by a balance between receptor depletion and renewal at the cell surface. **(A)** In untreated cells, the internalization and degradation of receptors is balanced by the synthesis and maturation of new receptors, and their renewal at the cell surface. Arrow size indicates relative rate. **(B)** In the presence of TGF-β, internalization and degradation is faster than renewal, so receptors become depleted from the cell surface. **(C)** In the presence of Activin, internalization and degradation are matched by renewal, so no depletion is seen. **(D)** In the presence of BMP4, the balance is transiently tipped towards internalization and degradation due to the up-regulation of SMAD6/7, depleting receptors from the cell surface. In the presence of longer durations of BMP4, SMAD6 and SMAD7 are down-regulated and internalization and degradation are again matched by renewal. Receptors re-accumulate at the cell surface.

Our mathematical modeling approach has suggested for the first time the importance of ligand affinity for receptors in shaping the signaling dynamics. We have shown that we can convert our mathematical model from simulating the refractory behavior observed for TGF-β to the non-refractory, integrated signaling behavior observed for Activin and BMP, by reducing the affinity of receptors for their ligand. This suggests that it is the high affinity that TGF-β that has for its receptors, (which is likely, at least in part, to be due to the cooperative interaction between TGF-β and the TGF-β type I and type II receptors (Hinck, 2012, Groppe et al., 2008)), that explains how TGF-β binding leads to a dramatic depletion in surface receptors, and the subsequent refractory behavior. In contrast, Activin, which binds its receptors with lower affinity, may not saturate the cell surface receptors, and thus does not cause obvious cell surface receptor depletion. In the case of BMP4, our experimental and modeling results indicate that it essentially functions like Activin, but the activity of the induced SMAD6/7 causes a transient depletion of receptors from the surface and a subsequent dip in PSMAD1/5 levels, giving the characteristic single oscillatory behavior.

The differences in surface receptor depletion seen in response to TGF-β, BMP4 and Activin also explains the differences in the integration of signaling observed over the first hour after stimulation. The constant presence of BMP and Activin receptors at the surface results in a continuous increase in receptor activation over the first hour, such that a longer duration of ligand exposure leads to more receptors being activated. Because the R-SMADs monitor receptor activity as a result of their nucleocytoplasmic shuttling, accumulation of activated receptors results in accumulation of phosphorylated R-SMADs (Schmierer et al., 2008). In the case of TGF-β, receptor activation is maximal after 5–10 min and does not continue to increase with time of ligand exposure.

### Distinct TGF-β family signaling dynamics may account for the different *in vivo* roles for these ligands

The differences in signaling dynamics that we have uncovered may account for the distinct roles these ligands play during embryonic development and tissue homeostasis. Activin, and the related ligand NODAL, as well as the BMPs, are well known to form gradients to pattern tissues, and are thought to act as morphogens (Wharton et al., 1993, Gurdon et al., 1994). Crucially, these ligands are all regulated by soluble extracellular ligand antagonists, such as Chordin or Noggin for BMPs, Follistatin for Activin, and Lefty1/2 for NODAL, among others (Brazil et al., 2015, Hedger and de Kretser, 2013, Schier, 2009). The formation of morphogen gradients requires cells to be sensitive to ligand levels at all times and both the BMP and NODAL gradients formed in early zebrafish embryos have been shown to be shaped by the action of ligand antagonists (Schier, 2009, Pomreinke et al., 2017, Ramel and Hill, 2013, van Boxtel et al., 2015, Zinski et al., 2017).

In contrast to Activin, NODAL and BMPs, TGF-β itself has never been shown to act in a gradient during embryonic development. The main roles of TGF-β during early stages of development are in facial morphogenesis (Dudas et al., 2006), heart valve formation (Mercado-Pimentel and Runyan, 2007) and in the development and maintenance of the vascular system (ten Dijke and Arthur, 2007), and graded ligand activity is not apparent in any of these processes. Furthermore, unlike Activin, NODAL and the BMPs, TGF-β has no known natural ligand antagonists. Like all the TGF-β family ligands, TGF-β is synthesized as a precursor, with a large prodomain and a C-terminal mature domain. The mature domain is then cleaved from the prodomain by proteases of the subtilisin-like pro-protein convertase (SPC) family (Miller and Hill, 2016). This pro-mature complex forms a latent complex with latent TGF-β binding proteins (LTBPs), and a further activation step is required to release mature TGF-β protein (reviewed in (Miller and Hill, 2016). Activin and BMPs are also secreted as pro-mature complexes, but their pro and mature domains are only weakly associated (Mi et al., 2015, Wang et al., 2016). It has been demonstrated for Activin that the pro and mature domains have a dissociation constant of ~ 5 nM and thus will be mostly dissociated at the concentrations required for full bioactivity (Wang et al., 2016). Thus, active TGF-β is only generated when and where it is required, while Activin and BMPs are essentially secreted as active ligands. We speculate that in the absence of any natural antagonists, the refractory behavior exhibited by TGF-β after stimulation may be a defence against deregulated signaling, such as occurs in cancer and fibrosis (Akhurst and Hata, 2012).

Morphogen gradients have been shown to be gradients, not just of ligand concentration, but also of time (Kutejova et al., 2009). In the current paradigm, both the amount and the duration of ligand exposure determines the fate of a cell in a gradient. For the Activin, NODAL and BMP pathways, where signaling receptors accumulate over time while ligand is present, the levels of PSMAD are proportional to signal duration and ligand dose. In contrast, a cell in a TGF-β gradient would be unable to measure the duration of its exposure to ligand, as almost signaling is initiated within the first few minutes. Moreover, a putative ligand antagonist would be unable to neutralize TGF-β, as most of the signaling occurs from internal compartments. Thus, TGF-β is regulated at the level of ligand production and release from the latency complex, and does not form signaling gradients.

### BMP exhibits an oscillatory behavior

We have demonstrated an oscillation in signaling downstream of BMP4 in multiple cell lines. This behavior depends on the transient upregulation of SMAD6 and SMAD7, which are required for the transient depletion of BMPR2 from the cell surface, that in turn correlates with the transient attenuation of signaling. The next step will be to investigate whether oscillations downstream of BMP signaling are observed in *in vivo* systems and what their function is. An attractive possibility is that they could be involved in periodically providing competence for cell fate decisions. It has been hypothesized that oscillatory behavior of both BMP and Notch signaling is required for vascular patterning, in particular, in sprouting angiogenesis, to determine the selection of tip versus stalk cells (Moya et al., 2012, Beets et al., 2013). This idea was based on the scattered expression of *Id1/2/3* (prominent BMP target genes) in the mouse angiogenic epithelium, which was postulated to reflect a snapshot of non-synchronized oscillatory gene expression. It will be very interesting in the future to directly monitor BMP signaling live in this system, to determine whether such oscillations occur.

## Materials and Methods

### Cell lines, and treatments

The human keratinocyte cell line, HaCaT, the human breast cancer line MDA-MB-231, the mouse fibroblast cell line NIH-3T3 and the mouse teratoma cell line P19 were used throughout this study. All cells were maintained in DMEM (Thermo Fisher Scientific), supplemented with 10% FCS. Ligands and reagents were used at the following concentrations: TGF-β (Peprotech), 2 ng/ml; BMP4 (Peprotech), 20 ng/ml; Activin A (PeproTech), 20 ng/ml; Noggin (PeproTech), 500 ng/ml; Follistatin (Sigma), 500 ng/ml; LDN-193189 (Gift from Paul Yu), 1 µM; SB-431542 (Tocris), 10 µM; Cycloheximide (Sigma), 20 µg/ml; Actinomycin D (Sigma) 1 µg/ml. The TGF-β neutralizing antibody, 1D11, and isotype-matched IgG1 monoclonal control antibody raised against Shigella toxin (13C4) were as described (Nam et al., 2008), and used at 30 µg/ml. All stimulations were performed in full serum. Where ligands or drugs were washed out, cells were washed three times with warm media. Whole cell extracts were prepared as previously described (Inman et al., 2002b). Where required, cell lysates were treated with PNGase F (New England Biosciences), 500 U per 100 µg of protein.

### Surface biotinylation and immunoblotting

Surface biotinylation assays were as previously described (Vizan et al., 2013). Immunoblotting were performed using standard techniques with the following antibodies: anti-PSMAD2 (Cell Signaling Technology, Cat. # 3108), anti-SMAD2/3 (BD Biosciences, Cat. # 610843), anti-PSMAD1/5 (Cat. # 13820), anti-SMAD1 (Invitrogen, Cat. # 38-5400), anti-ACVR1B (Abcam, Cat. # Ab133478), anti-ACVR2B (Aviva Systems Biology, Cat. # ARP45041, anti-BMPR2 (BD Biosciences, Cat. # 612292), anti-TGFBR1 (Santa Cruz, Cat. # sc-398), anti-TGFBR2 (Santa Cruz, Cat. # 17792), anti-TDGF1 (Cell Signaling Technlogy, Cat. # 2818), anti-MCM6 (Santa Cruz, Cat. # sc-9843), anti-Tubulin (Abcam, Cat. # Ab6160). Western blots were visualized on film or using an ImageQuant LAS 4000 mini (GE Healthcare) and quantified with ImageJ. For quantifications, densitometry measurements were normalized to loading controls and are shown relative to levels in cells stimulated with ligand for 1 hr, except where indicated.

### qPCR and siRNA knockdown

qRT-PCR was performed as previously described (Gronroos et al., 2012). Primer sequences are given in Supplementary file 1. For siRNA experiments, cells were plated, and 24 hr later transfected with 30 nM siRNA/3 µl RNAiMax (Thermo Fisher Scientific) for NIH-3T3 cells and P19 cells or 5nM siRNA/8 µl INTERFERin (PolyPlus) for MDA-MB-231s and 200 µl Opti-MEM (ThermoFisher Scientific) in fresh media. Volumes are given for a 6-well plate. Experiments were performed 72 hr after siRNA transfection. siRNAs were purchased from Dharmacon and sequences are given in Supplementary file 1. They were used as SMARTpools.

### Statistical analysis

Student’s t-tests were performed where appropriate using GraphPad Prism 7 software.

### Mathematical modeling

The mathematical models of Activin and BMP signaling are based on our previously published model of TGF-β signaling (Vizan et al., 2013), with the following key modifications.

Ligand binding to competent surface receptors is now treated as a reversible process. In the original TGF-β model, the dissociation rate of the ligand/receptor interaction was considered negligible compared to the activation of the receptor complex by the ligand, and ligand binding was treated as irreversible for simplicity. In the new model, this reaction is made reversible to allow modeling of different binding affinities of different ligands. An off-rate 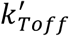 was thus introduced.

In addition, a negative feedback mechanism mediated by I-SMADs was included to model the behavior of cells in response to BMP4. I-SMADs were assumed to be synthesized in response to ligand, and to promote the degradation of signaling competent receptors, as well as the ligand-induced increase in degradation of active receptors.

I-SMADs are transcriptional targets of nuclear R-SMAD–SMAD4 complexes. Both I-SMAD RNA and protein were included to capture the time delay between ligand addition and I-SMAD expression. The two new equations for *I-SMAD* RNA and I-SMAD protein read:

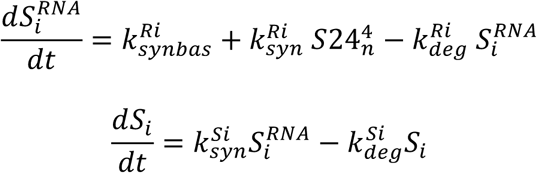

With these modifications, equations 2–5 from (Vizan et al., 2013) now read (new terms indicated in bold):

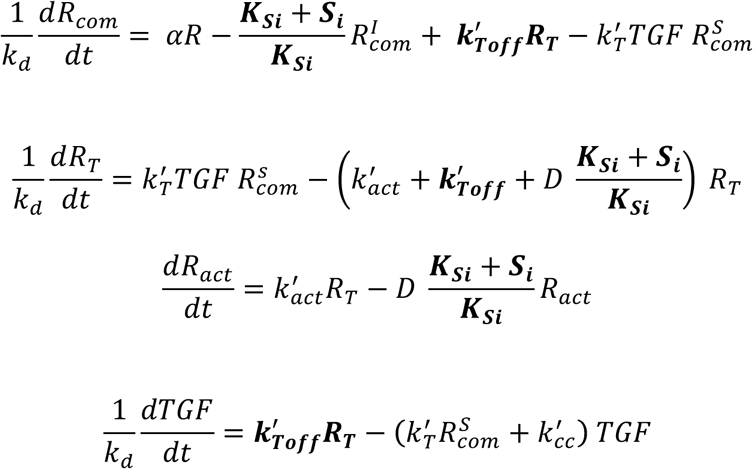

The following parameters were used to model the behavior of the I-SMADs.

| Parameter | Value |
| --- | --- |
| $k_{synbas}^{Ri}$ | 0 |
| $k_{syn}^{Ri}$ | 4 |
| $k_{deg}^{Ri}$ | 1 |
| $K_{Si}$ | 0.02 |
| $k_{syn}^{Si}$ | 1 |
| $k_{deg}^{Si}$ | 1 |

We have implemented these changes into a single model that can capture the dynamics of each ligand simply by changing the parameters in each case. The following parameters were changed to model each ligand, with key parameter changes indicated in bold:

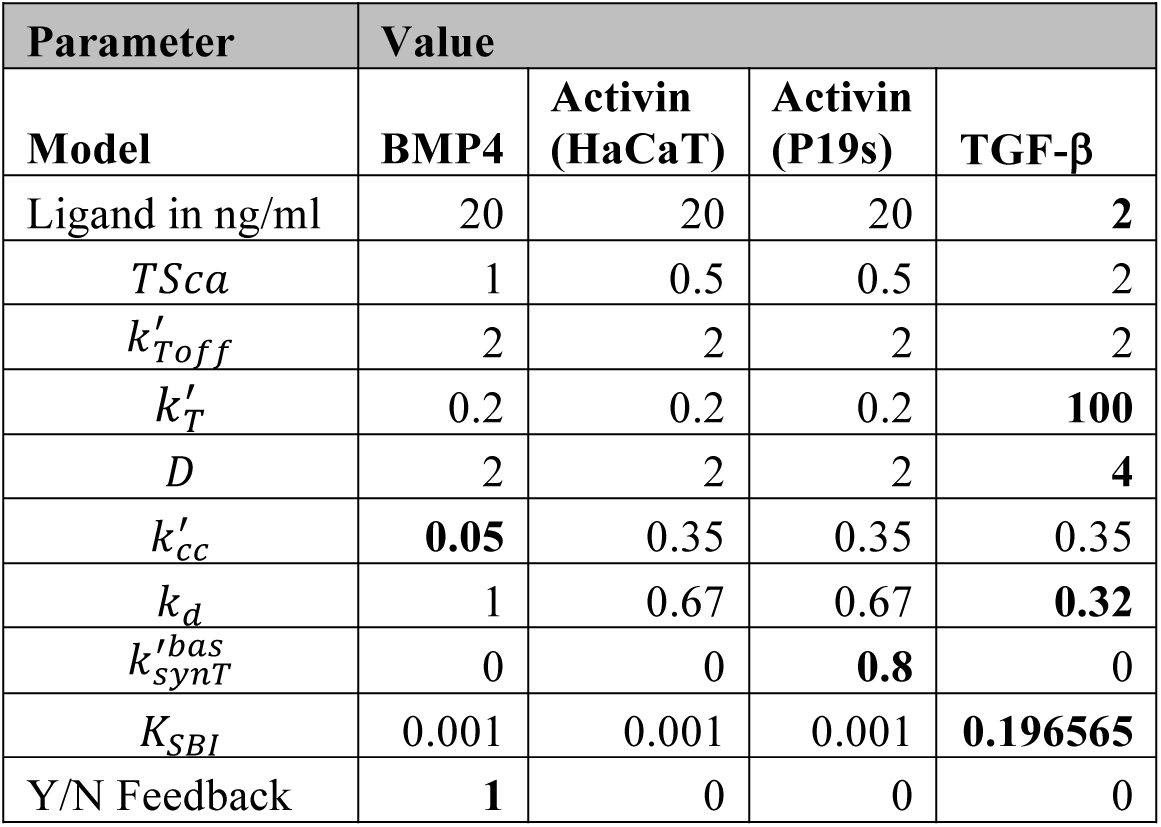

The key parameters changed are as follows:

- The on-rate of ligand to receptor binding, 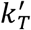, was chosen such that the dissociation constant of the ligand/receptor interaction, which is given by 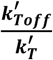 is very small for TGF-β (reflecting the high affinity of this ligand for its receptors), and is much larger for the other ligands. This is the only critical change necessary to alter the overall behavior of the model in response to each ligand.
- 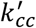 is the constitutive clearance of the ligand from the medium. Assuming that BMP4 is cleared from the medium at the same speed as the other ligands does not model the data well; it seems to be more persistent in the medium.
- 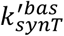 is the basal ligand production, which is required for modeling Activin dynamics in P19s, which secrete ligand in an autocrine fashion.
- *K*_*SBI*_is the dissociation constant of SB from the receptors.
- Y/N Feedback is a toggle switch that allows us to switch on and off I-SMAD production in response to ligand.

In addition, alterations to the following parameters were necessary to accurately capture the experimental data:

- *k*_*d*_ is the half-life of receptors in the absence of ligand.
- *D* is the ligand induced increase in degradation of active receptors
- *TSca* scales the relative amounts of ligand to receptor

The model was implemented in the freely available software packages COPASI (http://www.copasi.org) and XPP (http://www.math.pitt.edu/~bard/xpp/xpp.html). All simulations and parameter fitting were performed in COPASI (Hoops et al., 2006). The model has been deposited in the Biomodels database (Chelliah et al., 2015) and assigned the identifier *MODEL1810160001* and will be made publicly available after curation.

## Acknowledgments

We are grateful to the help we have received from Francis Crick Institute Cell services. We thank Paul Yu for the LDN-193189 and Lalage Wakefield for 1D11 and the isotype-matched control antibody. We thank all the members of the Hill lab and An Zwijsen for useful discussions and Anassuya Ramachandran for comments on the manuscript. This work was supported by the Francis Crick Institute, which receives its core funding from Cancer Research UK (FC001095), the UK Medical Research Council (FC001095), and the Wellcome Trust (FC001095).

## Author contributions

CSH and DSJM designed the study, DSJM performed all the experiments and BS performed the mathematical modeling. CSH and DSJM wrote the manuscript with input from BS.

---

**Figure 1 – figure supplement 1.**
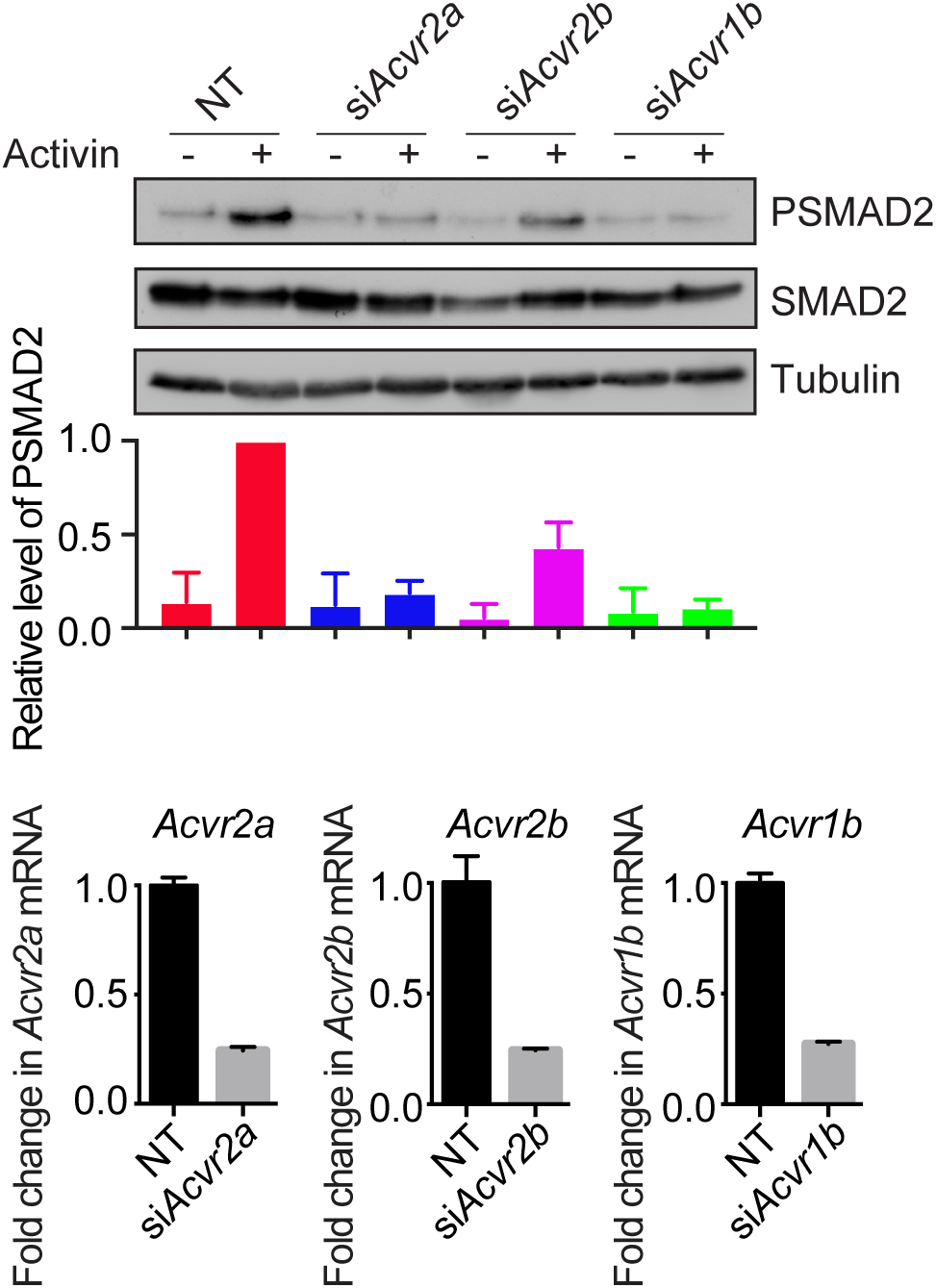
Characterization of Activin receptors in P19 cells. P19s were transfected with siRNAs against *Acvr2a, Acvr2b* or *Acvr1b*, then treated or not with Activin for 1 hr. Western blotting for PSMAD2, SMAD2 and Tubulin as a loading control was performed. Quantifications are the normalized means and SDs of densitometry measurements from two independent experiments. Below, the extent of knockdown was determined by qPCR. Shown are the normalized averages and SDs from two independent experiments, expressed as fold change in mRNA level relative to NT controls.

**Figure 1 – figure supplement 2.**
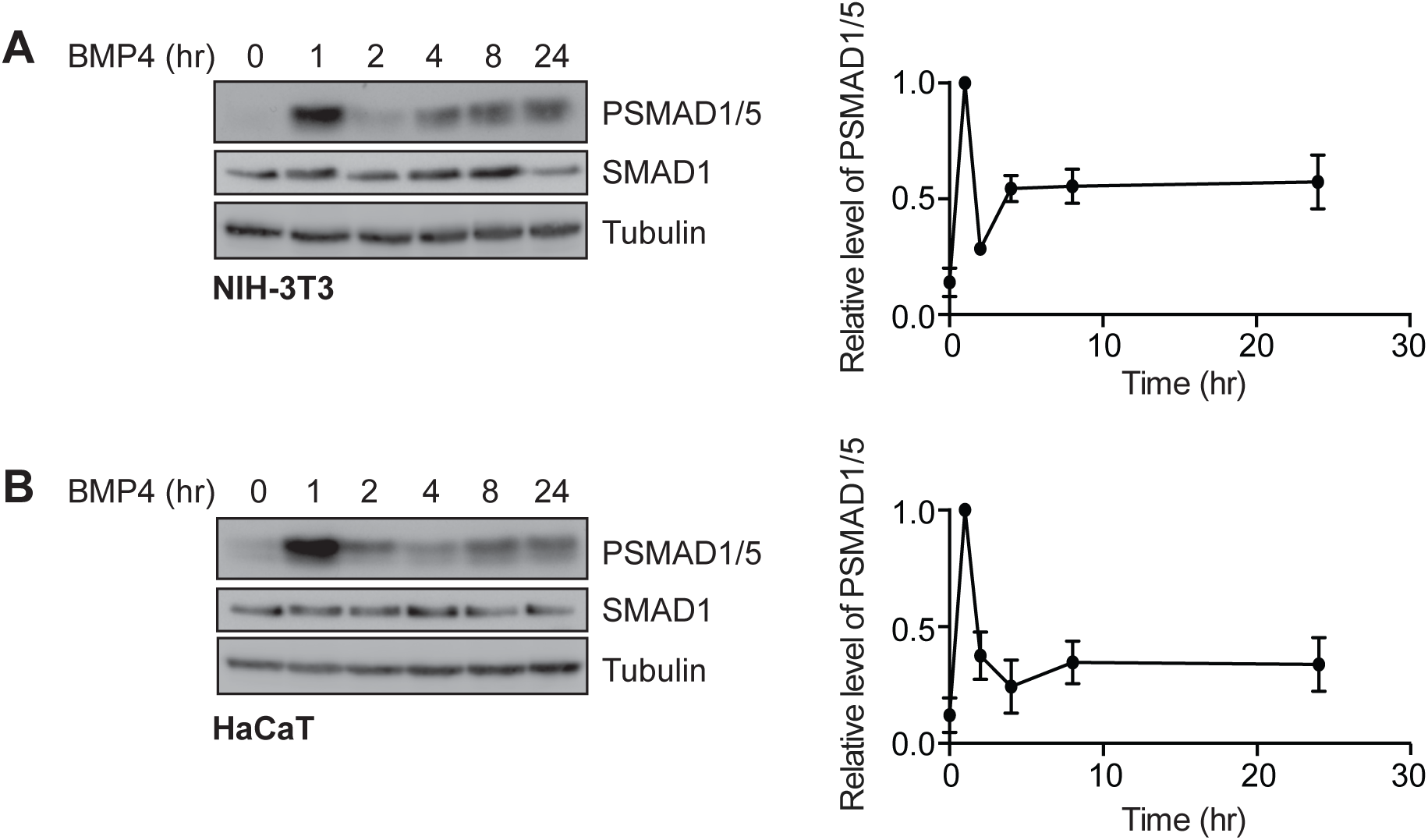
BMP4 exhibits oscillatory signaling in NIH-3T3 cells and in HaCaTs. **(A)** NIH-3T3s or **(B)** HaCaTs were treated with BMP4 for the times indicated. Western blotting for PSMAD1/5, SMAD1 and Tubulin as a loading control was performed. Quantifications are the normalized means and SDs of densitometry measurements from three independent experiments.

**Figure 5 – figure supplement 1.**
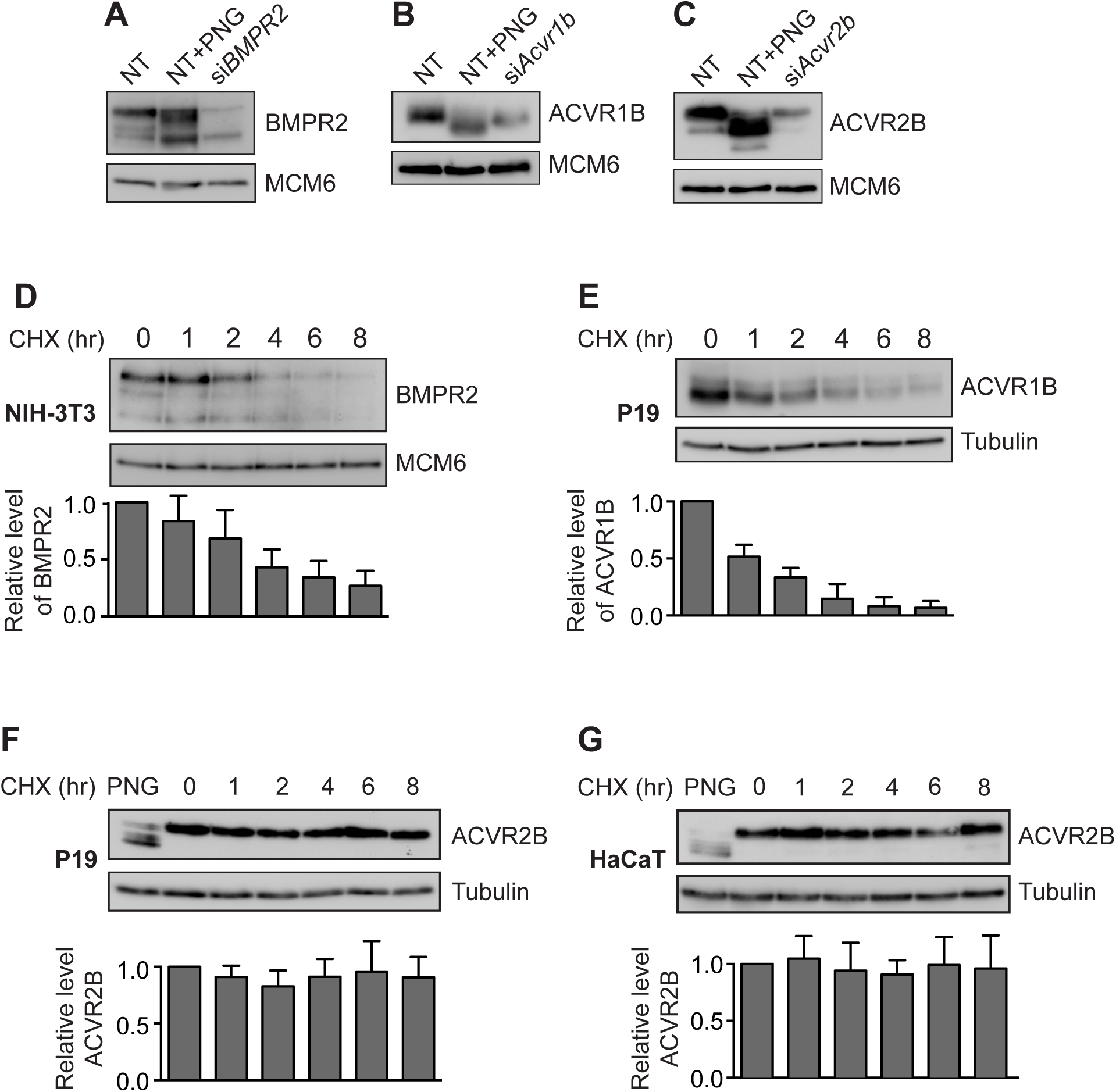
Characterization of receptor stabilities. **(A)** MDA-MB-231 cells were transfected with siRNAs against NT controls or BMPR2. Lysates were treated or not with PNGase (PNG). **(B)** P19 cells were transfected with siRNAs against NT controls or *Acvr1b*. Lysates were treated or not with PNGase (PNG). **(C)** P19 cells were transfected with siRNAs against NT controls or *Acvr2b*. Lysates were treated or not with PNGase (PNG). **(D-G)** NIH-3T3, P19 cells or HaCaTs as indicated were treated with cycloheximide (CHX) for the times indicated. Western blotting for BMPR2, ACVR1B, ACVR2B and Tubulin or MCM6 as a loading control was performed. In all cases, quantifications are the normalized means and SDs of densitometry measurements from three independent experiments relative to levels in untreated cells.

**Figure 6 – figure supplement 1.**
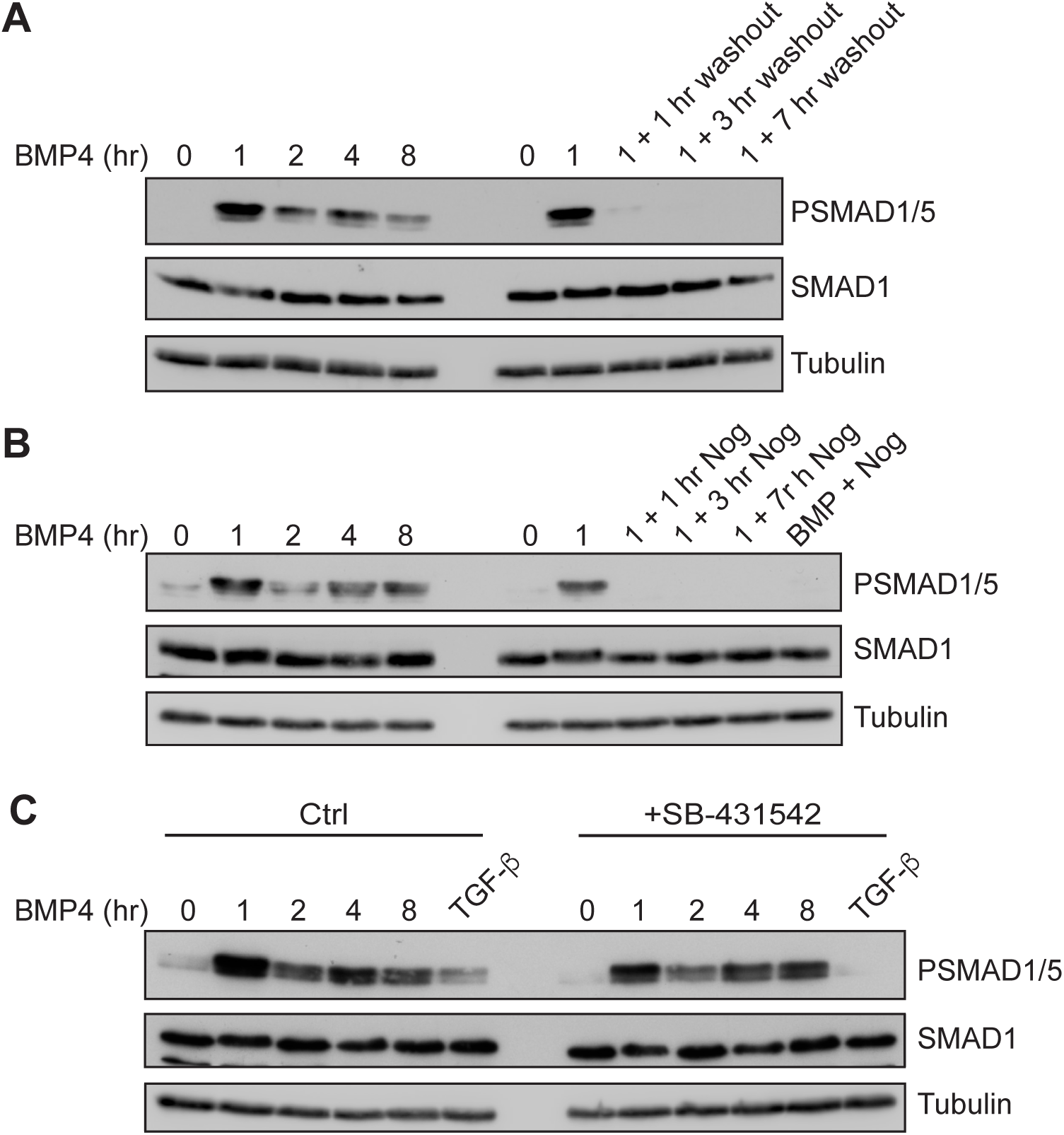
The BMP4 oscillation requires persistent exposure to BMP4 and is not mediated indirectly via SMAD2/3 signaling. **(A)** NIH-3T3 cells were treated with BMP4 for the times indicated, or for 1 hr with BMP4, before washout and incubation for the times indicated. **(B)** NIH-3T3s were treated for BMP4 for the times indicated, or for 1 hr with BMP4 before addition of Noggin for the times indicated. In the final lane, BMP4 and Noggin were added simultaneously and cells incubated for 1 hr. **(C)** NIH-3T3 cells were stimulated with BMP4 for the times indicated or with TGF-β for 1 hr, in the absence (Ctrl) or presence (+SB) of SB-431542. Western blotting for PSMAD1/5, SMAD1 or Tubulin as a loading control was performed. Representative blots from two independent experiments are shown.

**Figure 6 – figure supplement 2.**
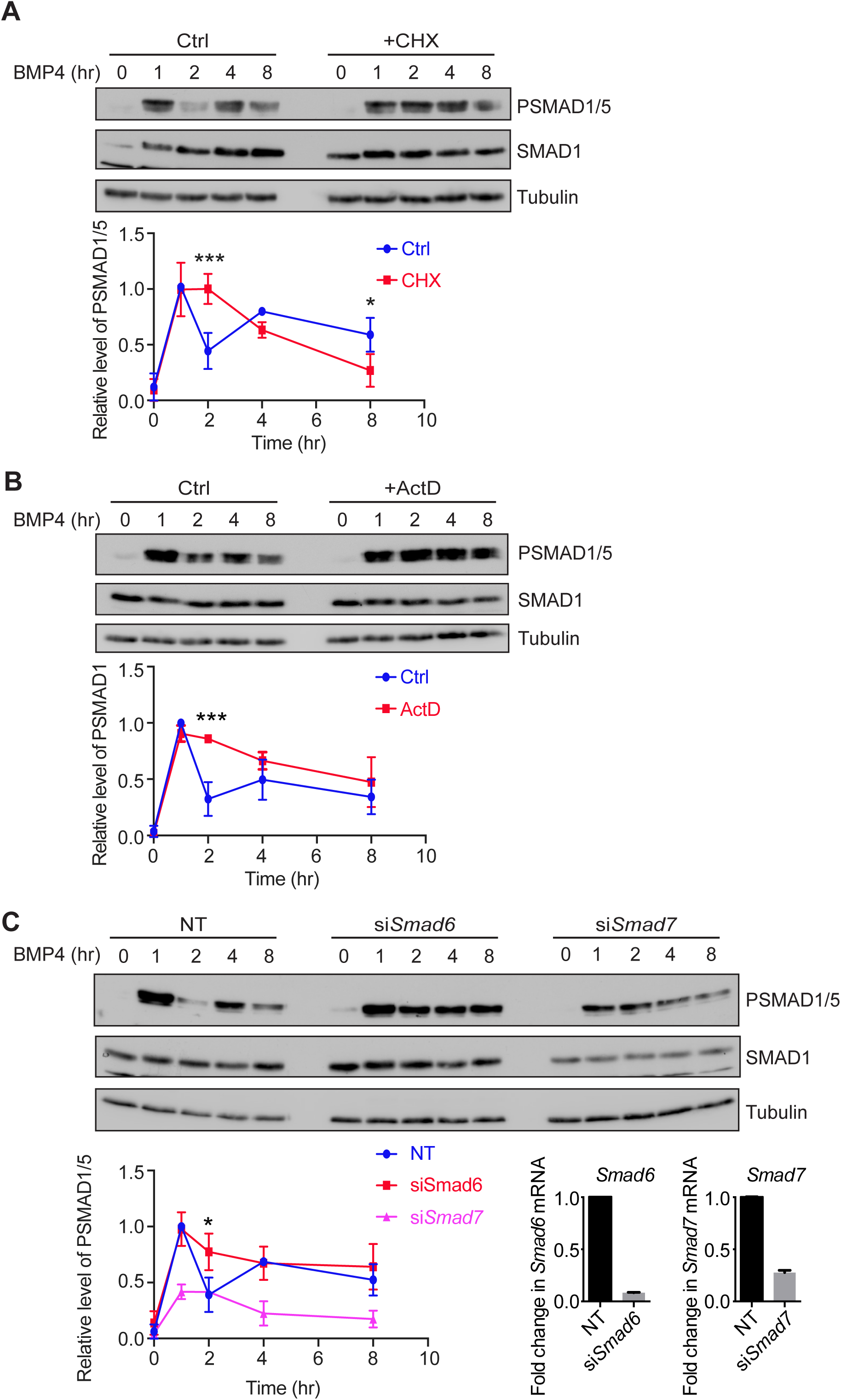
The BMP4 oscillation requires new protein synthesis. **(A)** NIH-3T3 cells were pre-treated or not with cycloheximide (CHX) for 5 mins, followed by BMP4 for the times indicated. **(B)** NIH-3T3 cells were pre-treated or not with Actinomycin D (Act D) for 5 mins, followed by BMP4 for the times indicated. In both cases, Western blotting for PSMAD1/5, SMAD1 and Tubulin as a loading control was performed. Quantifications are the normalized means and SDs of densitometry measurements from three independent experiments. *** indicates p<0.0005 **(C)** NIH-3T3 cells were transfected with siRNAs against NT controls, *Smad6* or *Smad7* and stimulated with BMP4 for the times indicated. Western blotting for PSMAD1/5, SMAD1 and Tubulin as a loading control was performed. Quantifications are the normalized averages and SDs of densitometry measurements from three independent experiments. * indicates p<0.05. Below right, the extent of knockdown was determined by qPCR. Shown are the normalized averages and SDs from two independent experiments, expressed as fold change in mRNA level relative to NT controls.

**Figure 6 – figure supplement 3.**
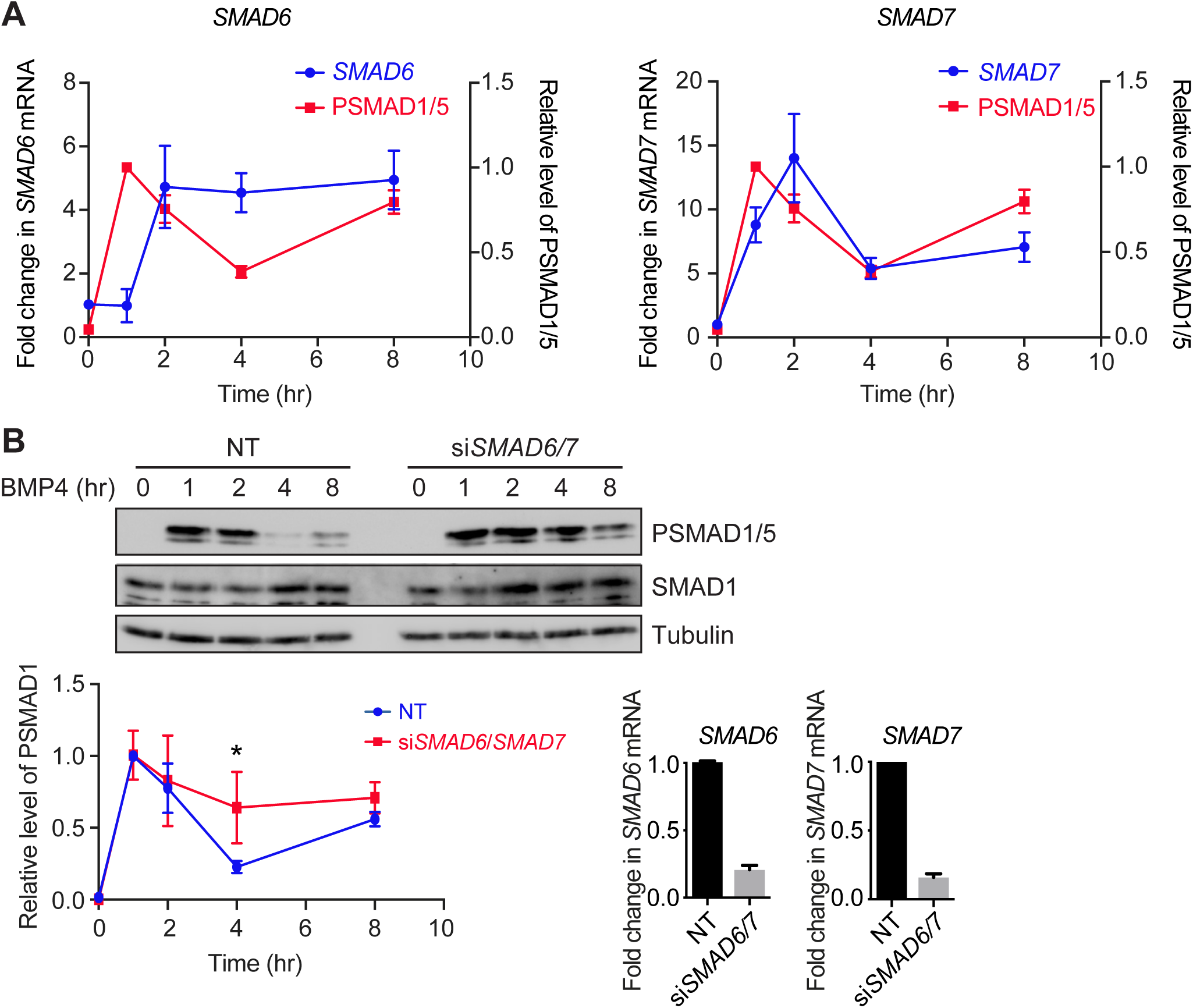
The BMP4 oscillation requires SMAD6/SMAD7 in MDA-MB-231 cells. **(A)** MDA-MB-231 cells were treated with BMP4 for the times indicated. Levels of *SMAD6* and *SMAD7* mRNA were assayed by qPCR. Shown are the normalized averages and SDs from three independent experiments, expressed as fold change in mRNA level relative to untreated cells. The PSMAD1/5 levels are from the data shown in Figure 1A. **(B)** MDA-MB-231 cells were transfected with non-targeting control siRNAs (NT) or siRNA SMARTpools targeting *SMAD6* and *SMAD7*, and were then treated with BMP4 for the times indicated. Western blotting for PSMAD1/5, SMAD1 and Tubulin was performed. Quantifications are the normalized means and SDs of densitometry measurements from three independent experiments. * indicates p<0.05. The extent of knockdown was determined by qPCR. Shown are the normalized means and SDs from three independent experiments, expressed as fold change in mRNA level relative to NT controls.

